# A Modular Chromosomal Passenger Complex Rewires Chromosome Segregation in *Plasmodium berghei*

**DOI:** 10.1101/2025.09.05.674402

**Authors:** Magali Roques, Chengyue Niu, Mathieu Brochet, Lorenzo Brusini

## Abstract

Faithful chromosome segregation relies on precise kinetochore–microtubule interactions and checkpoint surveillance, yet the molecular basis of these processes varies widely across eukaryotes and is only beginning to be defined in apicomplexan parasites. In the malaria parasite *Plasmodium berghei*, chromosome segregation is especially critical during transmission from host to mosquito: rapid mitoses generate male gametes, and subsequent meiosis in the zygote seeds the next generation of infection. Here, we identify Aurora-related kinase 1 (ARK1) as a central regulator of chromosome segregation in both mitotic and meiotic contexts. ARK1 localises to spindle poles, spindles, and kinetochores, and its depletion results in short and multipolar spindles, kinetochore misalignment, and failed chromosome partitioning. ARK1 forms a minimal Chromosomal Passenger Complex (CPC) with INCENP1 during male gametogenesis, but a more elaborate CPC with INCENP2, kinetochores, centromeric histones, and spindle assembly checkpoint proteins during meiosis. This stage-specific modularity allows *Plasmodium* to prioritise male gamete formation whilst safeguarding faithful chromosome inheritance during zygote development, ensuring parasite transmission to the mosquito. Our findings demonstrate that conserved CPC principles are rewired in *Plasmodium*, highlighting both the plasticity of eukaryotic checkpoint control and a potential vulnerability for blocking malaria transmission.

## Introduction

Faithful segregation of eukaryotic chromosomes during cell division depends on their attachment to the microtubule-based spindle via kinetochores (Cheeseman and Desai, 2008). At mitosis, duplicated chromosomes are segregated precisely once to each daughter, whereas the production of haploid gametes requires two consecutive meiotic divisions: homologous chromosomes are segregated in meiosis I, followed by separation of sister chromatids in meiosis II (Miller et al., 2012). Fidelity of chromosome segregation depends upon biorientation at metaphase, sister chromatids bound by microtubules emanating from opposing spindle poles (Foley and Kapoor, 2013). In animals and fungi, biorientation is assured through an error-correction process: incorrect kinetochore-microtubule attachments trigger a “wait” signal via the spindle assembly checkpoint (SAC), a surveillance system that delays the onset of anaphase until all chromosomes are correctly attached (Musacchio and Salmon, 2007). Correction of erroneous kinetochore-microtubule attachments is coordinated by antagonism between centromeric kinases and outer kinetochore phosphatases (Lampson and Cheeseman, 2011). Aurora B kinase as part of the chromosomal passenger complex (CPC)—along with INCENP, SURVIVIN, and BOREALIN—destabilizes improper attachments early in mitosis, activating the SAC (Liu et al., 2009). Upon biorientation, tension exerted across sister kinetochores is communicated through the action of PP1 and PP2A phosphatases, leading to checkpoint silencing and timely progression into anaphase (Lesage et al., 2011).

SAC control varies greatly across eukaryotes. Whilst the requirement for biorientation is widespread, SAC components and the mechanisms by which kinetochores communicate attachment status are not universally conserved (Meraldi et al., 2006; D’Archivio and Wickstead, 2016; Drinnenberg and Akiyoshi, 2017; van Hooff et al., 2017; Kops et al., 2020) . Within the phylum Apicomplexa—a group of intracellular parasites of considerable medical and veterinary relevance, including the malaria parasite *Plasmodium* and *Toxoplasma gondii*—plasticity in composition and organization of kinetochores across division modes provides a compelling case for the malleability of eukaryotic checkpoint control (Francia and Striepen, 2014; Hawkins et al., 2021; Brusini et al., 2022). Biorientation and an apparent delay in anaphase onset in response to deficient kinetochores, suggest the presence of at least some error-correction mechanisms within this group. However, SAC proteins (beyond a putative MAD1 homolog) and key kinases required for SAC recruitment, such as Monopolar Spindle 1 and Polo-Like, appear to be missing, questioning whether these parasites can support a robust error-correction response (van Hooff et al., 2017; Kops et al., 2020). Several kinases distantly related to Aurora and cyclin-dependent kinase families are required for DNA synthesis and mitosis in both *Plasmodium* and *Toxoplasma* (Reininger et al., 2011; Balestra et al., 2020; Hawkins et al., 2021; Zeeshan et al., 2023; Wyss et al., 2024). However, definitive roles and the presence of a CPC that coordinates correction of erroneous microtubule-kinetochore attachments during chromosome segregation remains unclear.

In this study, we uncover previously unrecognized, stage-specific differences in chromosome segregation and kinetochore organization during sexual development of *Plasmodium berghei.* These divisions are essential bottlenecks in the parasite lifecycle: failure to complete them faithfully directly compromises transmission from host-to-mosquito vector. We show that the Aurora-related kinase ARK1 associates with centromeric histones and kinetochores and is required for bipolar spindle formation, kinetochore alignment, and faithful segregation. Crucially, our results reveal a modular Chromosomal Passenger Complex (CPC) whose composition shifts between the mitosis of male gametogenesis and the meiosis of developing ookinetes. This flexibility reflects the distinct demands of division across host environments, enabling the parasite to balance speed with fidelity. Together, our findings illustrate how conserved CPC principles are rewired in *Plasmodium*, broadening our understanding of the plasticity of eukaryotic checkpoint control.

## Results

### The spindle MTOC partitions kinetochores during microgametogenesis

Sexual development in *Plasmodium berghei* is initiated by a change in conditions from host-to-mosquito following a blood-meal. Male gametocytes (microgametocytes) undergo three rounds of DNA replication and mitosis within minutes, producing eight flagellated male gametes. Male gametes fertilize female gametes (macrogametes), leading to meiosis in the zygote and subsequent differentiation into motile banana-shaped ookinetes.

Microgametogenesis does not appear to follow similar phases of mitosis as described in animal cells (Fig. 1A). In particular, microgametocyte mitosis lacks kinetochore alignment, linear structures formed by apicomplexan kinetochore proteins (AKiTs) as seen during asexual blood-stage mitosis (Fig. S1A) and highly reminiscent of biorientation at metaphase. Within the first three minutes post-activation, a spindle forms between tetrads of spindle pole MTOCs, each tethered to a basal body (Sinden et al., 1976, 1978), and together identified as a single bipartite focus by their dense NHS ester staining (Fig. 1A; panels i-ii “SM” & Rashpa and Brochet, 2022). Notably, the spindle contains more microtubules than would be expected based on the number of diploid (28) chromosomes. AKiT5 protein reveals that kinetochores (>28 AKiT5 foci, n=22) bind laterally to the spindle, with no clear evidence of end-on attachment during initial spindle formation. Subsequent movement of kinetochores is not driven by retraction of spindle microtubules, but by sequential partitioning of spindle pole MTOCs (Fig. 1A; panel iii-v “SM partition”) that halve the spindle 3 times during the following 6 minutes. One round of mitosis finally retracts spindle microtubules and segregates kinetochores culminating in eight kinetochore clusters, one destined for each flagellated male gamete.

**Figure 1.**
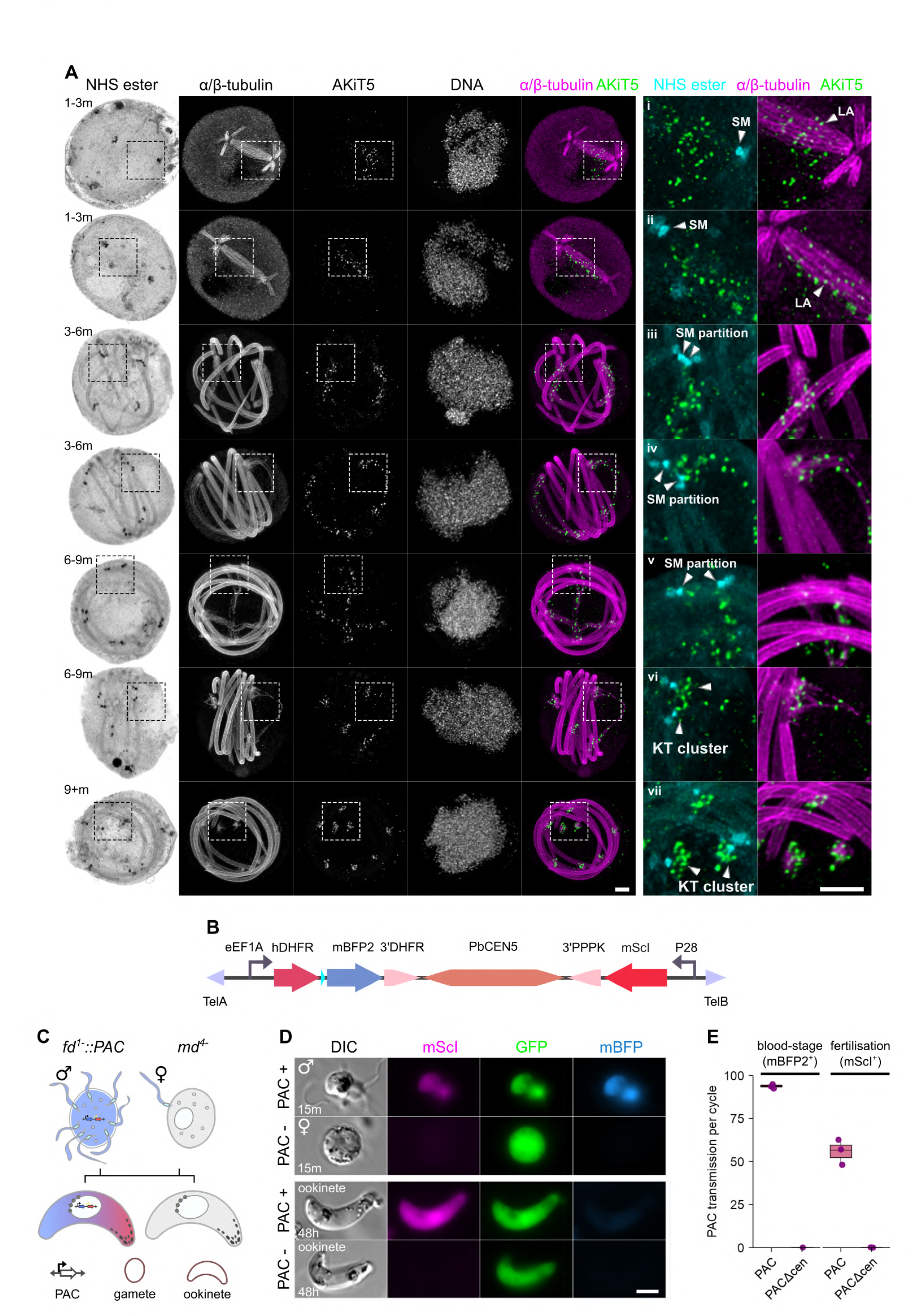
MTOC, spindle, kinetochore and chromosome dynamics during *Plasmodium* microgametogenesis. (A) U-ExM micrographs of *Plasmodium berghei* expressing the tagged kinetochore component AKiT5-mNeonGreen-3xHA (green) throughout microgametogenesis. Counter-staining of DNA (SYTOX), the spindle (α/β-tubulin) and spindle/basal body MTOCs (NHS ester) also shown. Minutes post-activation (m) shown on NHS ester panels. Representative panels displaying initial spindle formation between spindle pole MTOCs (SM) and lateral attachment (LA) of kinetochores (i-ii), partitioning of the spindle pole MTOCs (SM partition) and mitotic spindles (iii-v) and final segregation of kinetochores into clusters (KT clusters) at mitosis (vi-vii). Scale bar, 1 μm (non-expanded). (B) Schematic representation of engineered *Plasmodium* artificial chromosomes (PAC), encoding the centromeric region from chromosome 5, gene cassettes for the expression of mBFP2 and mScI and flanked by telomeric DNA. (C) Male-only (fd1-::PAC) and female-only (md^4-^) gametes were crossed and inheritance of PACs was monitored by fluorescence driven by eEF1α-driven mBFP2 (asexual stages) and P28-driven mScarlet-I fluorescence (ookinetes) by flow cytometry and fluorescence microscopy, respectively (D&E). Scale bar, 2 μm.

The lack of obvious kinetochore organization typically required for fidelity of chromosome segregation, at least in animal cells, raises the question as to whether chromosomes are faithfully inherited by gametes following microgametogenesis? To assess the fidelity of chromosome segregation during fertilisation, we took advantage of *Plasmodium* artificial chromosomes (PACs), linear DNA constructs encoding elements of the centromere and telomeres that facilitate faithful inheritance throughout the *P. berghei* lifecycle (Iwanaga et al., 2010). Here, these centromeres offer no selectable advantage to these cells but can be monitored due to the constitutive expression of the fusion of DHFR-mBFP2 driven by the eEF1α promoter, in addition to mScarlet-I (mScI) under the regulation of the female and ookinete promoter P28 (Fig. 1B). We engineered a male-only parasite line, *fd^1-^* (that constitutively expresses *GFP* driven by the eEF1α promoter from the *SSU* locus, (Sayers et al., 2024) to carry a PAC (*fd^1-^::PAC*) and compared the fidelity of PAC inheritance between intra-erythrocytic stages and during fertilisation, when crossed with untransformed female-only *md^4-^* gametes (Fig. 1C; and Fig. S1B). Fluorescence analysis identified mBFP2 fluorescence in a subset of blood-stage parasites following 14 intra-erythrocytic cycles equating to a minimum transmission fidelity of >93% per intra-erythrocytic cycle, compared to 0% of cells carrying a similar PAC, however lacking the centromere (Fig. 1D, E; and Fig. S1B, C, D, E; and Table. S1). In contrast, mScI fluorescence in developed ookinetes revealed fidelity of chromosome segregation to be far inferior during fertilisation, with PAC inheritance detected at between 55-60% in ookinetes 48 hours post-fertilisation. Surprisingly, mBFP2 fluorescence was not clearly detectable in ookinetes containing the PAC, despite clear GFP fluorescence, suggesting the gene contributed by the male gamete is transcriptionally silent, consistent with previous works showing *Plasmodium berghei* zygotes and ookinetes exhibit a maternal phenotype for constitutively expressed reporters (Ukegbu et al., 2015).

### Aurora kinase 1 associates with mitotic spindle poles, spindle and kinetochores during microgametogenesis

Our engineered PACs are inherited just over half the time following fertilisation in *P. berghei*. Importantly, this marked reduction in PAC inheritance compared to blood-stage division highlights a plasticity in chromosome stability or segregation mechanisms. We asked, what can the machinery between these modes of division tell us about fidelity of chromosome segregation in these parasites? Given the critical roles for Aurora kinases during mitosis in eukaryotic cells, we targeted an Aurora-related kinase called ARK1 that co-purifies with kinetochore components in gametocytes (Brusini et al., 2022), and for which the homolog in the human infecting species *Plasmodium falciparum* localises to spindle poles during intra-erythrocytic divisions (Reininger et al., 2011; Wyss et al., 2024).

To identify whether ARK1 also displays mitotic-like behavior during microgametogenesis, we localised protein in *P. berghei* alongside the outer-kinetochore marker NUF2-mScI and the inner-kinetochore marker AKiT5-mNeonGreen-3xHA (mNG-3xHA) (Brusini et al., 2022). Immunoblotting confirmed expression of the fusion protein at the expected mobility corresponding to the integration of the coding sequence for mBFP2-2xTy at the C-terminus of endogenous *ARK1* (Fig. S2A, B and C). Location of ARK1 fusion protein was diffuse in the nucleus throughout microgametogenesis (Fig. 2A), however accumulated to discrete foci and along spindle-like structures that tracked subsets of NUF2/AKiT5-marked kinetochores during mitosis. Signal for spindle/kinetochore-associated ARK1 puncta reduced to below detectability and nucleoplasm levels before exflagellation of male gametes. U-ExM confirmed ARK1 localisation at spindle pole MTOCs before activation (Fig. 2B) and subsequently along the tubulin microtubules and kinetochores during mitosis. ARK1 signal at spindles and kinetochores diminished shortly after completion of mitosis, approximately 12 minutes post-activation.

**Figure 2.**
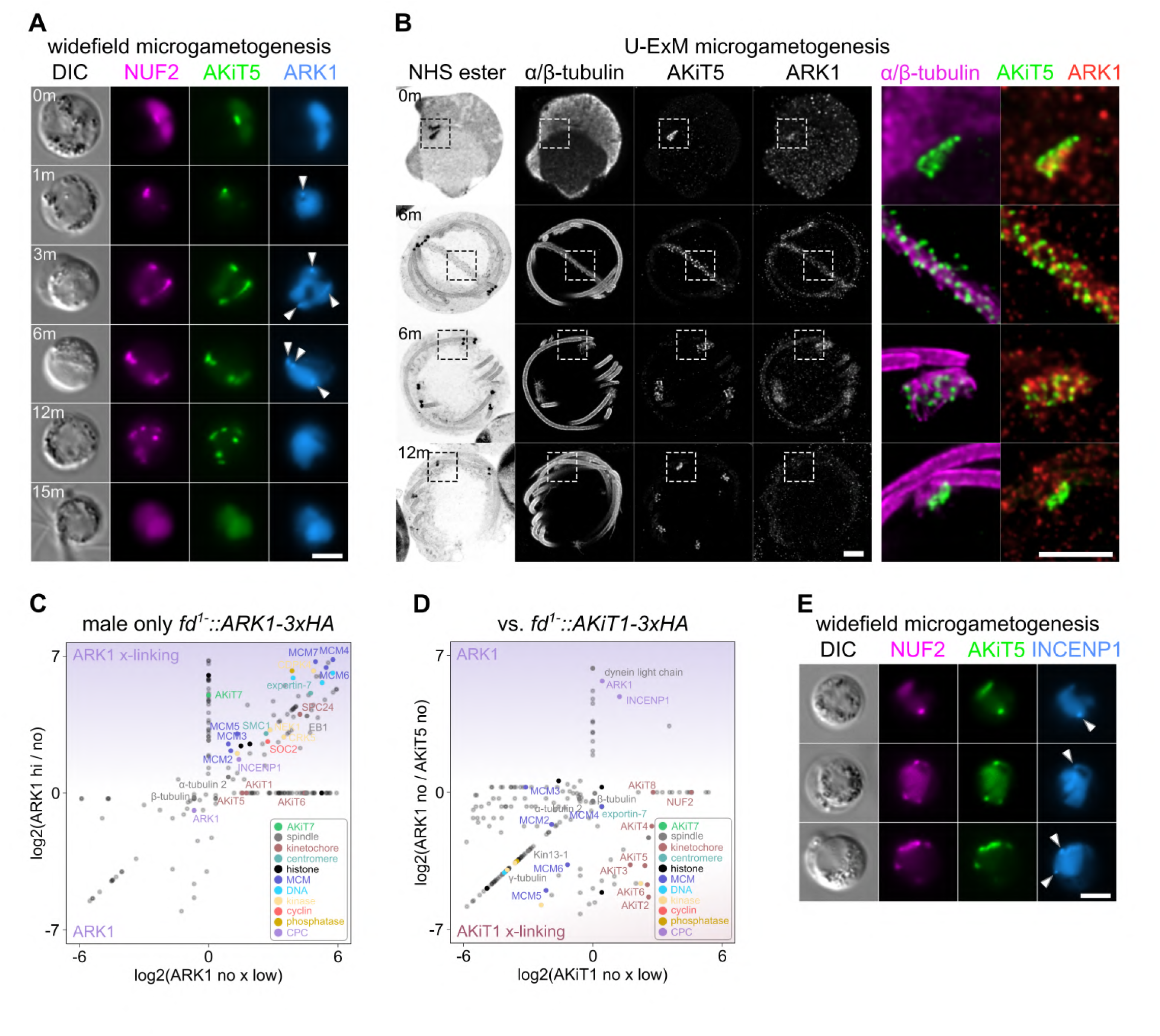
A CPC including ARK1 and INCENP1 localises to spindle poles and kinetochores in microgametocytes. (A) Micrographs of live native fluorescence in malaria parasites expressing tagged kinetochore components NUF2-mScI (magenta) and AKiT5-mNG-3xHA (green) and ARK1-mBFP2-2xTy during microgametogenesis. Differential interference contrast images (DIC) are also shown. Scale Bar, 2 μm. (B) Representative U-ExM micrographs displaying ARK1-mBFP2-2xTy in the nucleus during mitosis in microgametocytes, along with spindle pole MTOCs (NHS ester), mitotic spindles (α/β-tubulin) and kinetochores (AKiT5-mNG-3xHA). Scale Bar: 1 μm (non-expanded). (C&D) Relative enrichment of proteins identified by mass spectrometry following immunoprecipitation of ARK1-3xHA (magenta) under native and cross- linking conditions, and AKiT1-3xHA (burgundy), from male-only *fd^1-^*gametocytes. ARK1 and AKiT1-interacting proteins highlighted. Intensities of proteins not detected for either immunoprecipitation set to arbitrary minimum value. Intensities for all 302 proteins detected are presented in Table. S2.

To determine if ARK1 forms stable interactions with kinetochore or MTOC components, in addition to regulatory subunits analogous to the Chromosomal Passenger Complex (CPC) of Aurora B kinase (INCENP, SURVIVIN and BOREALIN), we used proximity cross-linking to identify interacting proteins and compared interactors to controls without cross-linking, as described for identification of AKiT proteins (Fig. S2D; Table. S2; Brusini et al., 2022) – the major difference here being that we affinity-purified protein from male-only *fd1^-^::ARK1-3xHA* gametocytes to distinguish from contaminating female interactions. In addition to α-/β-tubulin, a homolog of the inner-centromere protein called INCENP1 was among the most abundant interactors under both native and cross-linking conditions (Fig. 2C; Fig. S2E). Incremental cross-linking stabilised interactions between MCM helicase subunits and core histones. Spindle and kinetochore proteins, EB1 and SPC24 (Zeeshan et al., 2023, 2020), were enriched upon higher cross-linking conditions, in addition to components of the centromere, SMC1 and exportin-7 (Francia et al., 2020) and several kinases implicated in spindle function during microgametogenesis, including NEK1 (Zeeshan et al., 2024), CDPK4 (Billker et al., 2004) and the CDK-cyclin complex, CRK5- SOC2 (Balestra et al., 2020). The homolog of spindle assembly checkpoint protein, AKiT7/MAD1, was identified upon high cross-linking conditions alone. Comparative enrichment analysis pinpointed INCENP1 as the likely sole robust ARK1 partner under native conditions (Fig. 2D; Fig. S2F), while cross-linking of AKiT1 in kinetochores revealed the broader surrounding microtubule and chromatin landscape.

Consistent with biochemical interaction, localisation of INCENP1-mBFP2-2xTy during microgametogenesis closely mirrored that of ARK1-mBFP2-2xTy, present diffusely throughout the nucleoplasm and accumulating near NUF2/AKiT5-labeled kinetochores during mitosis (Fig. 2E, Fig. S2G).

### ARK1 and INCENP1 are required for bipolar spindle formation and kinetochore segregation during microgametogenesis

Our results indicate the presence of a reduced CPC, comprised of only ARK1-INCENP1, at spindle poles, spindle microtubules and kinetochores in microgametocytes. Given the conserved functions of CPCs in spindle assembly, kinetochore-microtubule attachments and chromosome partitioning across eukaryotes, we assessed the consequences of depletion on spindle integrity and kinetochore organization, by exchange of native 5’UTRs (Fig. S3A, B) to those of genes differentially regulated between sexual stage cells: *CLAG* (P_CLAG_), known to be lower in females relative to males (Sebastian et al., 2012), and *SKA2* (P_SKA_), a kinetochore component detected in female cells progressing to ookinetes but absent in male gametocytes (Brusini et al., 2022; Fig. S3C).

Depletion of ARK1 and INCENP1 (Fig. S3D), in particular under the regulation of P_SKA_, led to a severe reduction in microgametogenesis, with exflagellation rates falling to below 10% compared to parental lines 15 minutes post-activation (Fig. 3A). This effect was restored by ectopic PAC expression of full-length ARK1 but not ark1^K61R^, carrying the homologous mutation that renders human aurb^K106R^ inactive (Fig. 3B; Fig. S3E, F). Fluorescence analysis revealed a decrease in the number of NUF2/AKiT5-labeled kinetochore foci per microgametocyte (Fig. 3C), with fewer than 5% of P_SKA_-ark1 microgametocytes displaying 8 KT clusters 12 minutes post-activation compared to parental controls. U-ExM 3D-reconstructions (n=121; Table. S3; Fig. 3D; Video. S1) revealed no change in the number or morphology of spindle pole MTOCs (Fig. S3G, H; Video. S2), yet revealed a marked increase in cells harboring multipolar (Fig. 3E), short (Fig. 3F), or otherwise disorganized and defective spindles (Fig. 3G; Video. S3), unable to properly segregate kinetochores to daughter gametes (Fig. 3H). Notably, depletion of ARK1 also led to an over-amplification of outer MTOCs per microgametocyte, resulting in a marked increase in “free” outer MTOCs otherwise not connected to a spindle apparatus (Fig. S3I, J, K).

**Figure 3.**
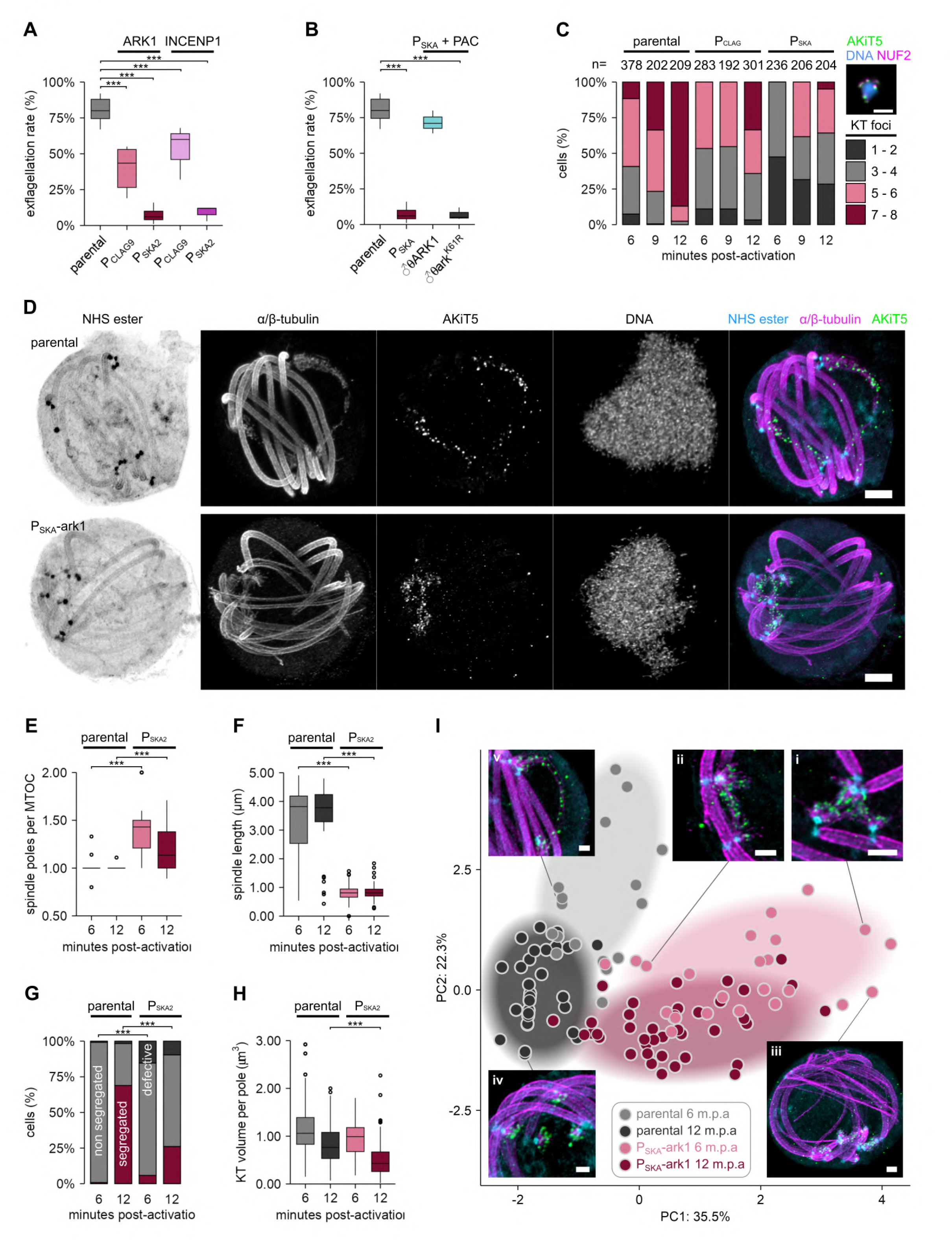
Depletion of ARK1 and INCENP1 impairs proper spindle and kinetochore dynamics during microgametogenesis. Morphological analyses of microgametocytes depleted for ARK1 and INCENP1 reveal reduced exflagellation compared to parental controls (A). This effect is rescued by ectopic male (♂) expression of full-length ARK1 but not by mutant ark1^K61R^ (B). Boxplots show the median (line), interquartile range (box), whiskers (1.5 × IQR), and outliers (points). Wilcoxon tests; ns ≥ 0.05; * < 0.05; ** < 0.01; *** < 0.001. (C) Quantification of kinetochore (KT) foci per nucleus in parental, P_CLAG_- and P_SKA_-ark1 lines at 6, 9, and 12 minutes post-activation. Stacked bar plots show the percentage of cells containing different numbers of KT foci (1–2, 3–4, 5–6, or 7–8), with total cell counts (n) indicated above each bar. Representative image (right) shows AKiT5 (green), NUF2 (magenta), and DNA (blue) staining. U-ExM of microgametocytes (D) revealed multipolar (E) and short mitotic spindles (F) in ARK1-depleted cells, unable to properly partition spindles (G) and segregate kinetochores (H) into daughter nuclei. MTOCs (NHS ester), mitotic spindles (α/β-tubulin), DNA (SYTOX) and kinetochores (AKiT5-mNG-3xHA) are shown. Scale Bar: 1 μm (non-expanded). Boxplots show the median (line), interquartile range (box), whiskers (1.5 × IQR), and outliers (points), Wilcoxon tests; ns ≥ 0.05; * < 0.05; ** < 0.01; *** < 0.001. Stacked barplots show the relative proportions of each group, χ² tests between mutant and parental at 6 and 12 min post-activation; ns ≥ 0.05; * < 0.05; ** < 0.01; *** < 0.001. (I) U-ExM measurements displayed in Principal Components (PC) 1 & 2 identifies clusters of cells corresponding to time points (6 vs 12 m.p.a) and cell line (parental vs P_SKA_-ark1). Representative micrograph panels displaying multipolar spindles (i), short spindles (ii), persistent spindles (iii), anaphase spindles (iv) and segregated kinetochore clusters (v), are also shown (α/β-tubulin; magenta, NHS ester; cyan, AKiT5; green). Scale bar, 0.5 μm.

Principal-component analysis (PCA) identified features strongly distinguishing ARK1-depletion at both 6 and 12 minutes post activation (Fig. 3I). The first two components encompass 58% of total variance (PC1 = 36 %, PC2 = 22 %). PC1 primarily captured variation in spindle polarity and structural integrity: cells with higher PC1 scores showed elevated spindle pole connections (Fig. 3I; panel i) and a greater proportion of short (Fig. 3I; panel ii) or defective spindles lacking kinetochores (Fig. 3I; panel iii), whereas lower PC1 scores marked cells with bipolar spindles and well-segregated kinetochores typical of parental lines (Fig. 3I; panel iv). PC2 reflected variation in mitotic progression and chromosome segregation, driven by average spindle length per cell and kinetochore volume per segregated cluster. This was particularly evident in parental lines, which showed a shift in PC2 from 6 to 12 minutes post activation, consistent with ongoing spindle elongation (Fig. 3I; panel v) and kinetochore maturation. In contrast, ARK1-depleted cells formed closer time point-independent clusters, indicating an initial arrest or persistent defect.

Taken together, ARK1 and INCENP1 form a minimal CPC that is indispensable for spindle polarity and kinetochore segregation during microgametogenesis. Their depletion results in multipolar and short spindles and defective kinetochore partitioning underscoring the essential role of this minimal CPC in parasite transmission.

### CPC function is critical to spindle elongation and end-on kinetochore attachments during meiosis, essential for mosquito transmission

In animal cells, antagonism between CPC kinase and PP1/PP2A corrects for erroneous spindle– microtubule attachments (Liu et al., 2009). In *P. berghei*, chromosome segregation at microgametogenesis appears unusually error-prone, suggesting a CPC incapable of supporting a robust error-correction response—despite critical roles in male gamete formation. To investigate whether changes in the CPC contribute to this vulnerability, we compared its function following transmission from host to mosquito vector.

Supporting the notion ARK1 and INCENP1 form core components of the CPC, depletion of either protein severely impacts oocyst infection in the mosquito midgut epithelium (Fig. 4A; top panel, x10 mag), remaining oocysts displaying necrotic features and aberrant kinetochore localisation (Fig. 4B; bottom panel, x63 mag). This defect is apparent within 8 days post-mosquito blood- meal, with a reduction in the number (Fig. 4B) and size of oocysts (Fig. 4C). Oocysts are further unable to properly form sporozoites, resulting in fewer migrating to salivary glands within 22 days post-mosquito blood-meal (Fig. 4D; all data in Table. S4).

**Figure 4.**
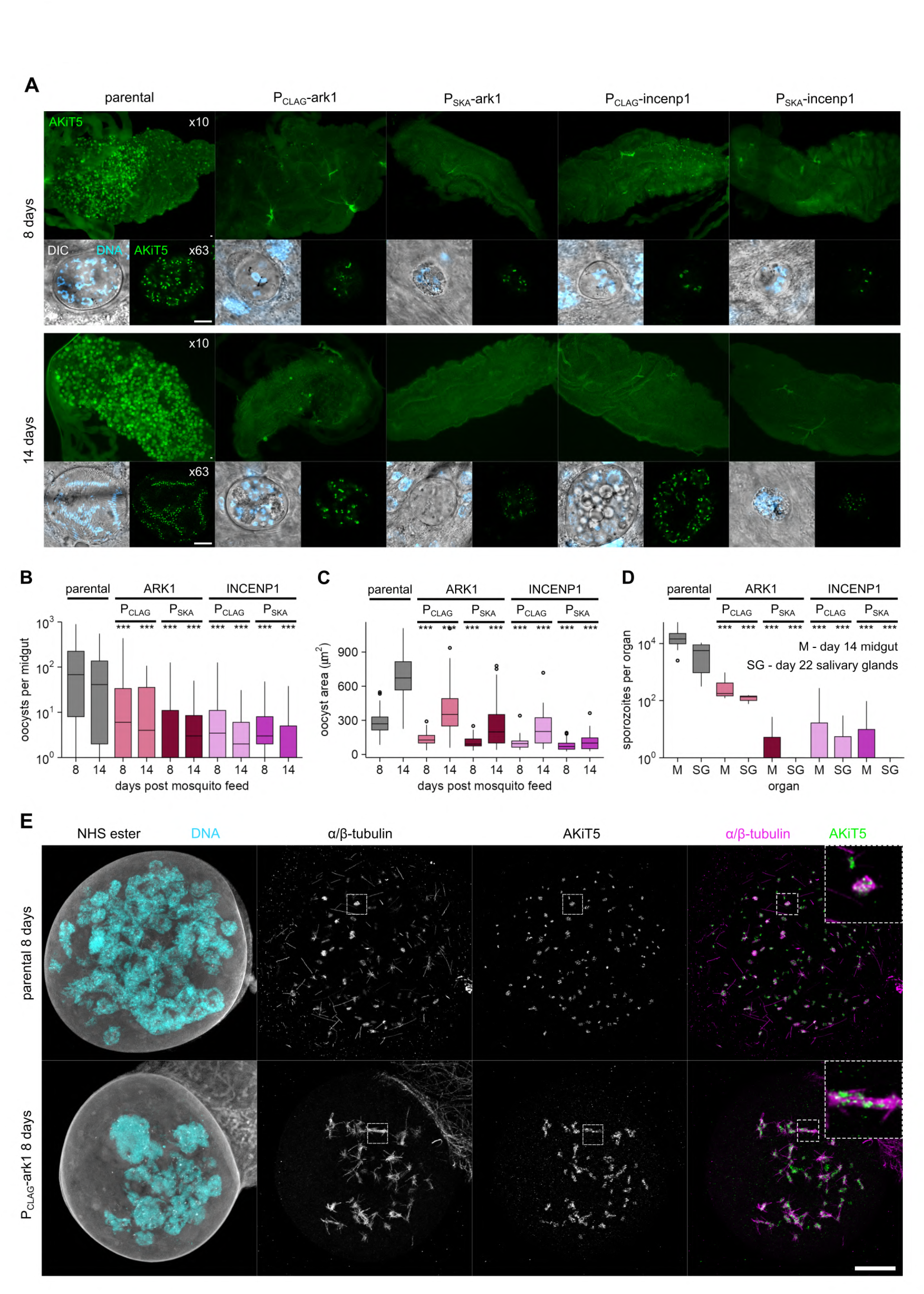
ARK1 and INCENP1 are essential for mosquito transmission. Live-imaging micrographs of mosquito midguts infected with oocysts expressing AKiT5-mNG-3xHA (green), from ARK1 and INCENP1-depleted cells and compared to parental controls, at both 8 and 14 days post-mosquito blood meal (top panels x10 widefield and bottom panels x63 confocal images shown, DNA stain Hoechst 33342 also shown in cyan). Scale bars, 10x; 100 μm, 63x; 20 μm. Total number (B) and size (C) of oocysts per midgut at 8 and 14 days post mosquito blood-meal, and migrating sporozoites per salivary glands at 21 days (D). Boxplots show the median (line), interquartile range (box), whiskers (1.5 × IQR), and outliers (points). Wilcoxon tests; ns ≥ 0.05; * < 0.05; ** < 0.01; *** < 0.001. (E) Representative U-ExM micrographs of P_CLAG_-ark1 and parental oocysts at 8 days of development. Protein labeling (NHS ester), DNA (SYTOX), spindle microtubules (α/β-tubulin) and kinetochores (AKiT5-mNG-3xHA) shown. Scale bar, 20 μm (non- expanded).

Notably, depletion of ARK1 or INCENP1 under the regulation of P_CLAG_, which caused only a mild reduction in microgametogenesis, had a strong effect during transmission. U-ExM further revealed ARK1-depleted oocysts with elongated mitotic spindles unable to properly partition kinetochores (Fig. 4E), however limitations in parasite numbers and constraints of imaging depth (working distance < 500 µm of 40x objective; Liffner et al., 2024) prevented further quantification.

To determine whether the vulnerability caused by depletion of CPC components ARK1 and INCENP1 during mosquito transmission arises from meiotic defects prior to colonization of the mosquito midgut epithelium, we tracked spindle and kinetochore dynamics in zygotes during ookinete development (Fig. 5A). Meiosis in *Plasmodium* deviates from most eukaryotes by occurring post-fertilisation, generating four distinct haploid genomes within a single nucleus. By 4 h post-fertilisation, the spindle MTOC duplicates and spindle microtubules emanate into the nucleoplasm (Fig. 5A; panels i&ii). AKiT5-marked kinetochores are unbound and only partially assembled, lacking outer kinetochore components of the NDC80-NUF2 complex (Zeeshan et al., 2020; Brusini et al., 2022). In contrast to microgametogenesis, following initial capture, kinetochores migrate along a bipolar meiotic spindle at metaphase to align centrally and establish end-on attachments to microtubules from opposing spindle poles (Fig. 5A; panel iii). Whilst ultrastructural studies have identified synaptonemal complexes between opposing kinetochore pairs (Sinden and Hartley, 1985; Sinden et al., 1985), these structures are not defined by NHS ester staining. Noteworthy is the presence of 3 spindle MTOCs and monopolar spindles in a subset of cells prior to metaphase alignment (Fig. 5A; panel iv*), indicating MTOC duplication is not dependent upon completion of the first chromosome segregation. By 12 h post-fertilisation, kinetochores segregate into separate clusters corresponding to anaphase (Fig. 5A; panel iv), coinciding with spindle collapse into a tight bundle during elongation. Two additional rounds of MTOC duplication and kinetochore segregation (Fig. 5A; panels v&vi), without apparent intervening metaphase alignments, reflect the sequential reductive divisions necessary for the migration of the nucleus (Fig. 5A; panel vii) along with 4 haploid kinetochore clusters into the body of mature ookinetes following 24 hours of development (Fig. 5A; panel viii).

**Figure 5.**
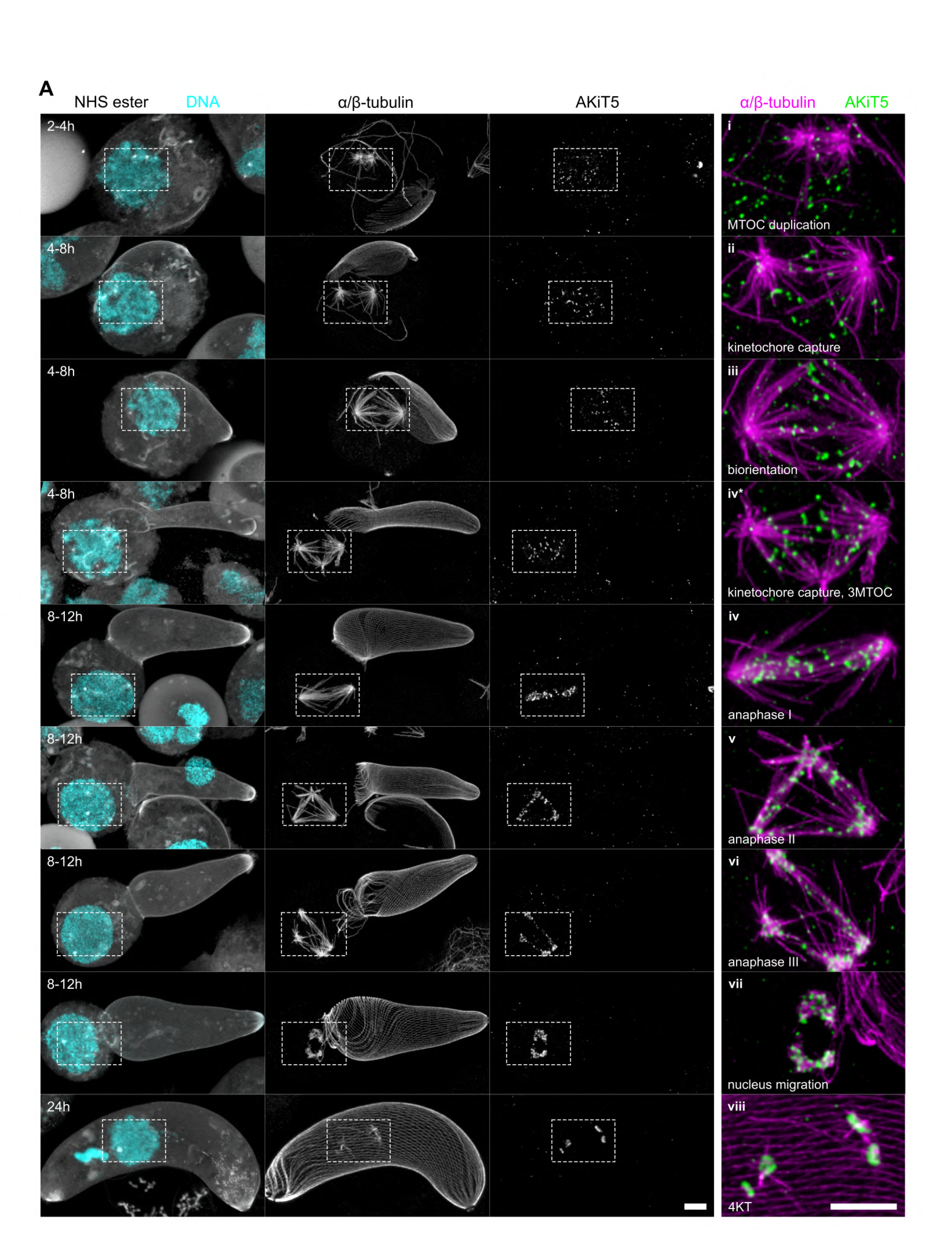
Spindle and kinetochore dynamics throughout *Plasmodium* meiosis. (A) U-ExM micrographs of *Plasmodium berghei* expressing the tagged kinetochore component AKiT5- mNeonGreen-3xHA (green) throughout meiosis. Counter-staining of DNA (SYTOX), microtubules (α/β-tubulin) and spindle/basal body MTOCs (NHS ester) also shown. Representative panels displaying MTOC duplication (i), initial kinetochore capture (ii), formation of bipolar spindle and biorientation of kinetochores (iii), second MTOC duplication prior to anaphase I seen in subset of cells (iv*), anaphase I-III (iv-vi), migration of the nucleus and kinetochores into the cell body (vii), and subsequent formation of four haploid kinetochore clusters in fully developed ookinetes (viii). Scale bar, 1 μm (non-expanded).

Consistent with a requirement during meiosis, ARK1-depleted cells fail to properly partition their kinetochores and display only 2 KT clusters in mature ookinetes—despite maintaining their characteristic banana-shaped morphology (Fig. 6A&B). Restoration of full-length ARK1 ectopically expressed from engineered PACs during microgametogenesis (Fig. 6C, Fig. S4A, “♂mSc”) confirmed this reduction originates in the zygote: only crosses where the male contributed functional ARK1 produced zygotes with four well-segregated KT clusters. In contrast, zygotes expressing kinase-dead ark1^K61R^ failed to restore proper segregation (Fig. 6C, Fig. S4B, “♂ARK1 + ♀ARK1/ark1^K61R^”).

**Figure 6.**
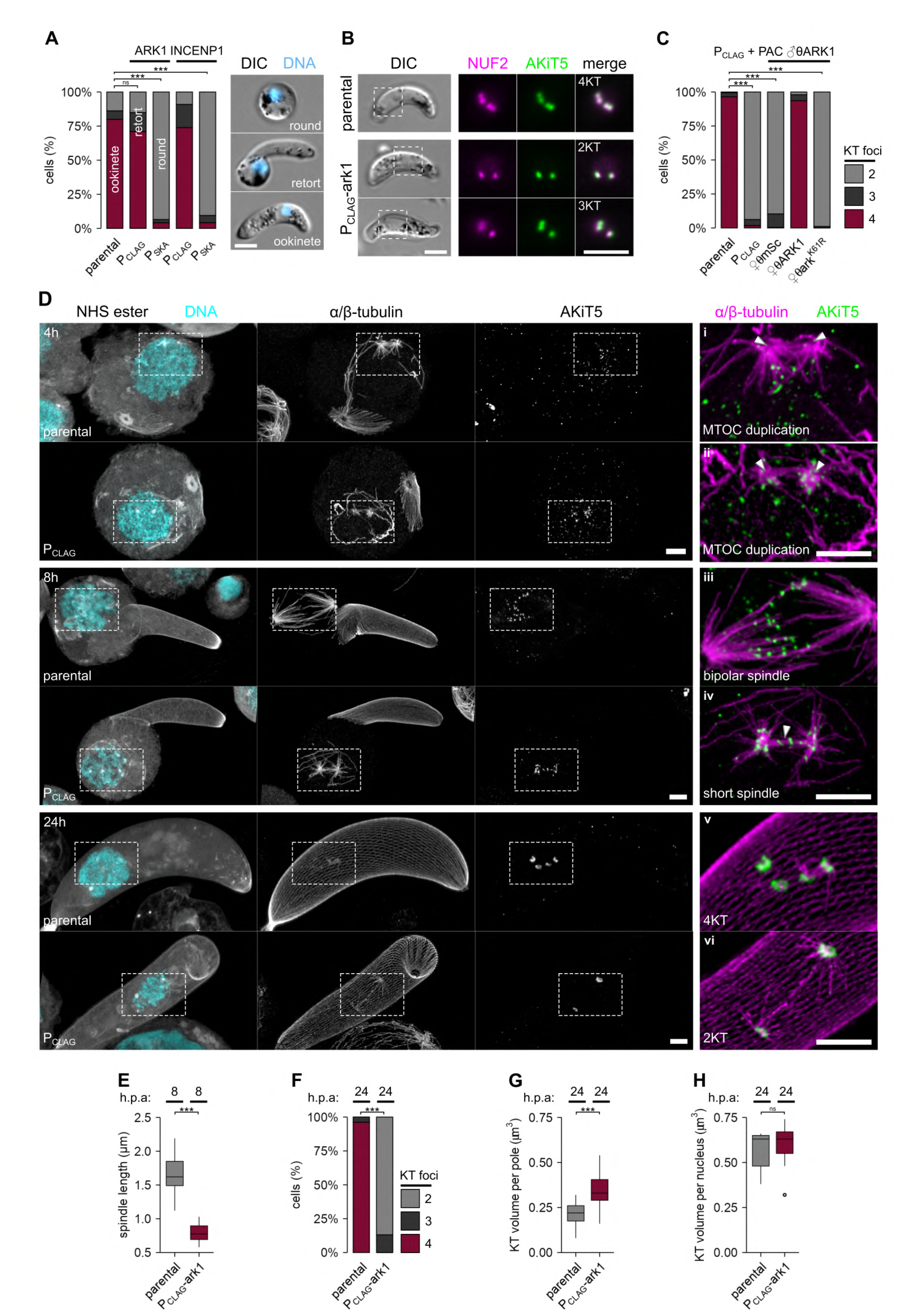
ARK1 depletion impairs meiotic spindle formation and kinetochore biorientation. (A) Morphological analyses of the products of fertilisation (female gametocytes and zygotes appear “round”, and developing ookinetes are either a “retort” or fully developed “ookinete”) in cells depleted for ARK1 or INCENP1 and compared to parental controls. Counter-staining of DNA (Hoechst 33342) also shown. Scale bar, 2 μm. (B & C) Depletion of ARK1 during ookinete development reduces the number of NUF2-mSc (magenta) and AKiT5-mNG-3xHA (green) labeled kinetochore foci in ookinetes compared to parental controls. This effect is rescued by ectopic male (♂) and female (♀) expression of full-length ARK1 but not by mutant ark1^K61R^. Stacked barplots show the relative proportions of each group, χ² tests between mutant and parental at 6 and 12 min post-activation; ns ≥ 0.05; * < 0.05; ** < 0.01; *** < 0.001. Scale bar, 2 μm. (D) Representative U-ExM micrographs of developing ookinetes depleted for ARK1 at 4, 8 and 24h post-fertilisation, revealing no clear effect of spindle pole duplication (α/β-tubulin; magenta) and nuclear morphology (SYTOX; cyan), however bipolar spindle length (E) and kinetochore attachment and biorientation (AKiT5-mNG-3xHA; green) are severely impaired, leading to abnormal numbers of kinetochore clusters in ookinetes (F). NHS ester counter-stain also shown in cyan. Scale bar, 1 μm. (G&H) Volume of kinetochore clusters per spindle pole and per nucleus, respectively. Boxplots show the median (line), interquartile range (box), whiskers (1.5 × IQR), and outliers (points), Wilcoxon tests; ns ≥ 0.05; * < 0.05; ** < 0.01; *** < 0.001. Stacked barplots show the relative proportions of each group, χ² tests between mutant and parental at 6 and 12 min post-activation; ns ≥ 0.05; * < 0.05; ** < 0.01; *** < 0.001.

U-ExM 3D reconstructions (Fig. 6D. Table. S5. n=85) revealed that although spindle MTOCs duplicate (Fig. 6D; panel i, ii), ARK1-depleted cells do not properly form a bipolar spindle, producing instead a short spindle (Fig. 6D; panel iii, iv. Fig. 6E; Video. S4, S5) unable to attach a full set of kinetochores to microtubules emanating from opposing spindle poles. This defect leads to unequal kinetochore partitioning (Fig. 6D; panel v, vi. Fig. 6F) and a complete block in meiosis, with no additional MTOC duplication required for tetraploid sets of kinetochores to form four distinct clusters in ookinetes following 24 hours of development (Fig. 6G, H; Video. S6, S7).

### A modular CPC interacts with the spindle, kinetochores and centromeres during meiosis

Our high resolution tracking of ARK1 function on spindle/kinetochore dynamics has revealed striking differences in chromosome segregation between mitosis and meiosis at mosquito transmission. Critically, ARK1 is required in meiosis for the formation of a bipolar spindle, proper kinetochore alignment at metaphase and equal partitioning of kinetochores, and depletion blocks the subsequent MTOC duplication required for the formation of haploid genomes, effects not seen following depletion of ARK1 during microgametogenesis mitosis. It is therefore possible that the CPC supports a more robust kinetochore – microtubule interaction and checkpoint signaling at meiosis compared to mitosis. To explore whether ARK1 behaviour may further suggest its meiotic capabilities, we similarly localised mBFP2 fusion protein alongside our markers for the inner and outer kinetochore. Location of ARK1 fusion protein was diffuse throughout ookinetes during development (Fig. 7A). In contrast to AKiT5 foci present in the nucleus shortly following fertilisation, ARK1 first accumulates as two distinct foci at the nuclear periphery following 2 h of development, consistent with MTOC duplication and spindle nucleation. By 8h of development, these foci relocate to opposing nuclear poles and along the bipolar meiotic spindle, concomitant with NUF2 and AKiT5 at kinetochores. Whereas kinetochore signals reveal two successive rounds of asynchronous duplication and migration, ultimately residing as four puncta at the nuclear periphery in mature ookinetes, ARK1 foci rapidly reduce to below detectability and cytoplasmic levels by 10 h of development. U-ExM confirmed ARK1 localisation at spindle MTOCs following duplication (Fig. 7B), in addition to discrete foci along bipolar spindle microtubules, many overlapping with both AKiT5 and NUF2 labeled kinetochores at meiosis.

**Figure 7.**
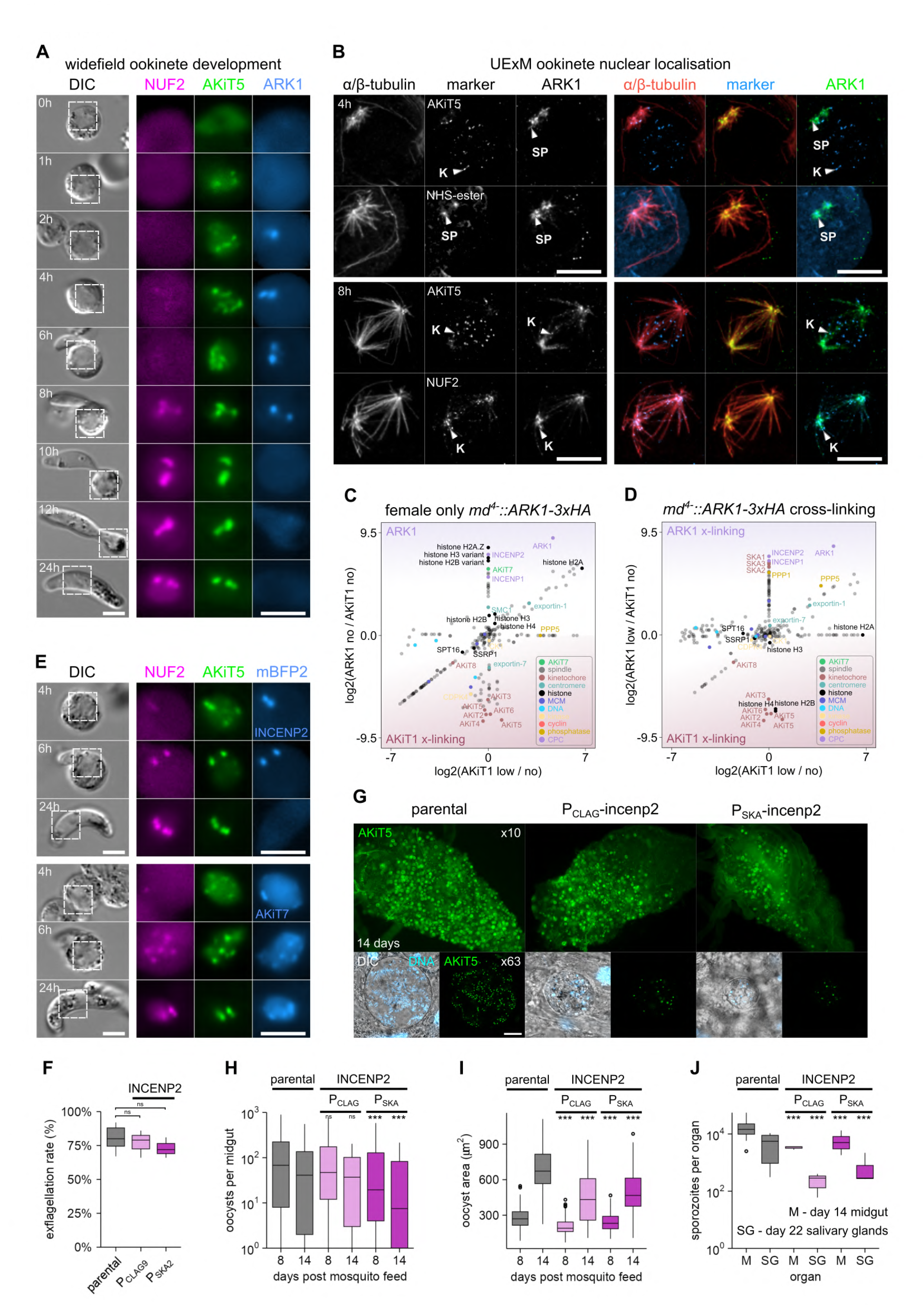
A CPC including INCENP2 interacts with the spindle and kinetochores during meiosis. (A) Micrographs of live native fluorescence in *P. berghei* developing ookinetes expressing tagged kinetochore components NUF2-mScI (magenta) and AKiT5-mNG-3xHA (green) and ARK1-mBFP2-2xTy (blue). Differential interference contrast images are also shown. Hours post-activation of gametocytes; “h”. Scale bar, 2 μm. (B) Representative U-ExM micrographs displaying ARK1-mBFP2-2xTy in the nucleus during meiosis, along with mitotic spindles (α/β-tubulin) and inner (AKiT5-mNG-3xHA) and outer (NUF2-mScI) kinetochore components. Hours post-activation of gametocytes; “h”. Scale Bar: 1 μm (non-expanded). (C&D) Relative enrichment of proteins identified by mass spectrometry following immunoprecipitation of ARK1-3xHA (magenta) under native and cross-linking conditions, and AKiT1-3xHA (burgundy) from female-only *md^4^*^-^ *P. berghei* 8 hours post-fertilisation. ARK1 and AKiT1-interacting proteins highlighted. Intensities of proteins not detected for either immunoprecipitation set to arbitrary minimum value. Intensities for all 324 proteins detected are presented in Table. S6. (E) Micrographs of live native fluorescence in *P. berghei* developing ookinetes expressing tagged kinetochore components NUF2-mScI (magenta) and AKiT5-mNG-3xHA (green) and either INCENP2-mBFP2-2xTy or AKiT7-mBFP2-2xTy (blue). Differential interference contrast images are also shown. Hours post-activation of gametocytes; “h”. Scale bar, 2 μm. Morphological analyses of microgametocytes depleted for INCENP2 reveal no clear reduction in exflagellation compared to parental controls (F). However live-imaging micrographs of mosquito midguts infected with oocysts expressing AKiT5-mNG-3xHA (green) from INCENP2-depleted cells reveal a marked reduction in infection (G; top panels x10 widefield and bottom panels x63 confocal images shown, DNA stain Hoechst 33342 also shown in cyan. Scale bars, 10x; 100 μm, 63x; 20 μm). Total number (H) and size (I) of oocysts per midgut at 8 and 14 days post mosquito blood-meal, and migrating sporozoites per salivary glands at 21 days (J). Boxplots show the median (line), interquartile range (box), whiskers (1.5 × IQR), and outliers (points). Wilcoxon tests; ns ≥ 0.05; * < 0.05; ** < 0.01; *** < 0.001.

To compare the composition of the CPC between mitosis and meiosis, we similarly immunoprecipitated ARK1 protein, however this time from female-only *md^4-^::ARK1-3xHA* zygotes purified 8 hours post-fertilisation by *fd^1-^* males, to enrich for interactions specifically during meiosis (Fig. 7C, D; Table. S6). Under native meiotic conditions, ARK1 co-purified not only with INCENP1, but along with a second homolog of the inner-centromere-protein INCENP2 among the most abundant interactors. Additional prominent stable interactions included the homolog of the SAC component AKiT7/MAD1 and several chromatin factors—histone variants H2A.Z, H3v, and H2Bv. Limited cross-linking of ARK1 stabilised interactions with the kinetochore–microtubule stabilizers SKA1-3 and PP1, factors which reinforce end-on kinetochore attachments and promote SAC silencing in eukaryotic cells (Pinsky et al., 2009; Vanoosthuyse and Hardwick, 2009).

To validate the sensitivity of our sex- and stage-specific strategy, and to contrast CPC composition between male and female interactomes during gametogenesis, we also performed ARK1 immunoprecipitations from mixed male and female *P. berghei 2.34* gametocyte populations 6 minutes post-activation (Fig. S5A; Table S6). In these mixed samples, ARK1 co-purified with both INCENP1 and INCENP2, as well as the SAC component AKiT7/MAD1. Importantly, similar INCENP2 or AKiT7/MAD1 interactions were not detected in our male-only ARK1 interactomes, demonstrating that these interactions arise exclusively from female cells. This observation highlights a key limitation of previous analyses (including our own), which examined ARK1 interactors in bulk gametocyte preparations: female-specific CPC components were inadvertently attributed to male mitotic complexes in earlier bulk gametocyte studies, whereas our sex- separated approach now resolves CPC composition with stage-specific precision.

Consistent with biochemical interaction, localisation of INCENP2-mBFP2-2xTy closely mirrored that of ARK1-mBFP2-2xTy during ookinete development (Fig. 7E), present as spindle pole-like foci that rapidly became undetectable following completion of meiosis. Conversely, AKiT7- mBFP2-2xTy resided primarily at the nuclear periphery and localised to kinetochores specifically between 6-8 h post-fertilization, together consistent with INCENP2 as the only likely additional component of the CPC during ookinete development.

Supporting the notion INCENP2 is a crucial component of the CPC specifically at meiosis, depletion has little effect on microgametogenesis (Fig. 7F; Fig. S5B, C), whereas oocyst development within the mosquito midgut is severely impaired (Fig. 7G, H). This effect is apparent within 8 days post-mosquito blood-meal, INCENP2-depleted oocysts are unable to grow to proper size (Fig. 7I), resulting in fewer sporozoites per oocyst and migrating to the salivary glands at 14 and 22 days post-mosquito blood-meal, respectively (Fig. 7J).

Taken together, INCENP2 expands the CPC specifically during meiosis, where it supports robust end-on kinetochore–microtubule attachments and oocyst development, highlighting a stage- specific vulnerability in parasite transmission.

## Discussion

ARK1 is a kinase we show is responsible for the precise orchestration of chromosome segregation, in both mitotic and meiotic contexts. It localises to spindles and kinetochores, forming part of a CPC essential for establishing bioriented kinetochore-microtubule attachments. But what do our observations for the CPC in *Plasmodium* mean within the broader eukaryotic kinetochore context? The discovery of kinetochore components and their biorientation at metaphase, in addition to the faithful transmission of PACs throughout the lifecycle, have challenged the traditional view that *Plasmodium* spp. possess a reductive, idiosyncratic machinery incapable of high-fidelity chromosome segregation (Arnot and Gull, 1998; Morrissette and Sibley, 2002; Iwanaga et al., 2010; Kops et al., 2020; Brusini et al., 2022). Nevertheless, *Plasmodium* kinetochores are comparatively streamlined relative to those in animals and fungi, containing fewer than ∼10 identified components across CCAN (AKiT9-11) and KMN (AKiT1-6) and missing key SAC proteins (except AKiT7/MAD1) and SAC-associated kinases (e.g., MPS1, PLK). Furthermore, our findings that ARK1 associates exclusively with INCENP to form the CPC are consistent with the apparent loss of SURVIVIN and BOREALIN from apicomplexan genomes (van Hooff et al., 2017; Komaki et al., 2022), and together point towards the substantial re-wiring of *Plasmodium* chromosome segregation machinery.

*Plasmodium* ARKs are Aurora kinases, a family with ubiquitous roles in spindle assembly, kinetochore function and cytokinesis. Evolutionary refinements have subfunctionalized paralogs in most modern-day eukaryotes, reflected not only by the limited phylogenetic signal linking Aurora families, even among animals (Hochegger et al., 2013), but by functional mosaicism in highly derived systems. In animals and plants, pairs of Aurora kinases (e.g. AURKB/C in mammals and AtAUR1/2 in *Arabidopsis*), overlap in function towards spindle assembly, CPC function and meiosis, while a third paralog remains less functionally conserved. Conversely, budding yeast employs a single Aurora kinase, IPL1, to control all Aurora-dependent processes, whereas the euglenozoan *Trypanosoma brucei* encodes three Aurora kinases, and a CPC formed by TbAUKB- INCENP-CPC2 that ensures accurate kinetochore-microtubule attachments during biorientation. *Plasmodium* ARKs represent a comparable example. ARK1 is required for bipolar spindle formation (AURKA-like), associates with the CPC (AURKB-like), and is required for proper meiotic progression (AURKC-like). ARK2 also exhibits spindle-associated behaviors reminiscent of AURKA and AURKB (Zeeshan et al., 2023), with ARK3 less characterized.

Genuine novelty in apicomplexan parasites appears in the presence of more recent duplications within core kinetochore and CPC sets, as previously described for the CCAN component CENP-C (AKiT9/10) and which we show here for INCENP. Phylogenetic patterns suggest this event occurred before the diversification of most modern apicomplexans, giving rise to the two INCENPs seen in Hematozoa and Coccidia and coinciding with the loss of SURVIVIN and BOREALIN (Berry et al., 2018; Komaki et al., 2022). The presence of two INCENPs within the *Plasmodium* CPC offers a modular system that can alternate between minimal and more stringent segregation machinery depending on developmental stage (Fig. 8). In particular, during sexual development this modularity permits a romantic love affair that “marries in haste, repents at leisure”, prioritizing a male-first programme that favours gamete output over maximal fidelity of chromosome inheritance. Chromosome segregation at microgametogenesis appears chaotic; kinetochores do not visibly congress or bi-orient, resulting in gametes that remain fertile despite incomplete genome integrity. ARK1 forms stable interactions exclusively with INCENP1 in microgametocytes. The absence of the SKA-C (Brusini et al., 2022) and apparent dispensability for INCENP2 correlate with the lack of clear metaphase structures that would typically be required to ensure faithful PAC transmission. Furthermore, ARK1 function during microgametogenesis suggests its primary roles may not be in spindle assembly checkpoint surveillance, consistent with observed multipolarity and short spindles seen in microgametocytes upon depletion. Notably, ARK1-depleted males also show over-amplified and ‘free’ outer MTOCs that are no longer connected to an intranuclear spindle, indicative of MTOC-spindle decoupling during microgametogenesis. This mirrors recent observations in *P. falciparum* asexual blood-stages, where disruption of NDC80-C disconnects the MTOC from the nucleus and perturbs spindle organisation, underscoring a functional coupling between the outer kinetochore and the MTOC (Li et al., 2024). Together with our recovery of EB1 and SPC24 as *bona fide* ARK1-proximal interactors, ARK1 activity likely also modulates outer-kinetochore NDC80-C function during microgametogenesis.

**Figure 8.**
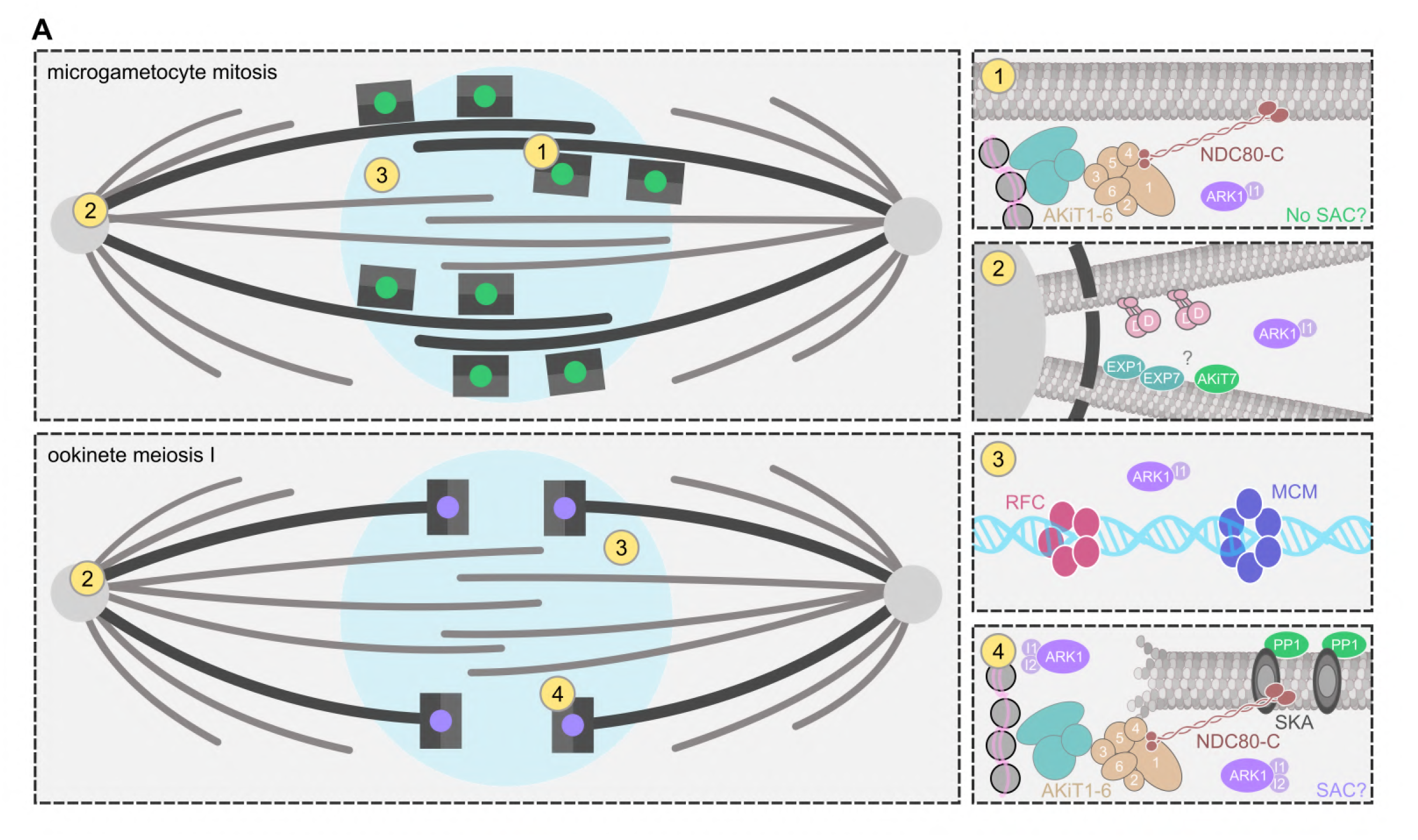
A modular CPC promotes spindle integrity and lateral-to-end-on kinetochore attachment, safeguarding faithful chromosome segregation in *P. berghei*. (A) Models for *P. berghei* chromosome segregation mechanisms. (1) Absence of INCENP2, SKA-C and kinetochore interactions with spindle assembly checkpoint proteins favor kinetochores forming lateral attachments to the spindle during mitosis in microgametocytes. ARK1 localises to spindle poles and along the spindle in microgametocytes where its presence is critical for spindle elongation and polarity (2). (3) Pools of ARK1 are also present proximal to DNA-interacting proteins of the MCM and origin of replication complexes. (4) At meiosis, additional interactions between ARK1 and INCENP2, the SKA complex and spindle checkpoint proteins promote robust end-on kinetochore attachments to the meiotic spindle.

Following fertilisation, zygotes restore order. Kinetochores align along the spindle midplane during meiosis I, and further MTOC duplication is delayed in the absence of proper chromosome segregation, together consistent with a surveillance mechanism and transmission-fidelity checkpoint that safeguards against the propagation of aneuploidy. ARK1 interacts with both INCENP1 and INCENP2 during meiosis, configurations required for CPC proximity to the spindle and kinetochore-associated SKA complex (SKA-C). During meiosis, ARK1 localises closer to the outer than to the inner kinetochore, further raising the possibility that ARK1 also directly influences outer kinetochore function to stabilise end-on kinetochore attachments to the spindle. Identification of PPP1 alongside SKA-C and ARK1 during meiosis strongly suggests the presence of a tension-sensing biorientation checkpoint. In animal cells, PPP1 antagonizes AURKB- mediated phosphorylation of SKA-C, stabilizing kinetochore-microtubule attachments and permitting anaphase onset under proper tension conditions—it is tempting to speculate that a similar relationship between ARK1-PPP1 senses tension across apicomplexan centromeres. Interestingly, ARK1 also forms stable interactions with histone variants H2A.Z, H3v, and H2Bv under native (i.e. non-crosslinking) conditions, whose presence likely reflects CPC docking onto specialized nucleosomes at the centromere (Fachinetti et al., 2013), in addition to the SAC component AKiT7/MAD1. It is quite possible therefore that INCENP2, rather than INCENP1, is the primary CPC subunit enabling ARK1 function in promoting chromosome biorientation. This requirement for INCENP2 within the CPC is also likely during asexual intra-erythrocytic stage divisions, given the clear kinetochore biorientation and localisation of ARK1 to subsets of mitotic nuclei in *P. falciparum.* Understanding the rationale behind these distinct CPC pools will further clarify the extent of SAC rewiring in *Plasmodium*.

A notable strength of our study is that we performed immunoprecipitations from genetically separated male-only and female-only parasite lines, using marker-free single-sex *fd^1-^* and *md^4-^ P. berghei* lines, resources recently developed by Sayers et al., that enable sex-resolved phenotyping and downstream biochemical applications (Sayers et al., 2024). This strategy removes the confounding influence of mixed gametocyte populations and provides stage-specific resolution of ARK1 interaction partners during mitosis or meiosis. Previous works, including our own proteomic surveys, have captured composite or transient associations, as seen with our identification of SKA-C (Brusini et al., 2022). The ability to disentangle male and female interactomes here offers a clearer view of CPC composition and rewiring across distinct developmental stages. Notably, our finding that AKiT7/MAD1 is recovered as a stable interactor primarily in female gametocytes, and localises to kinetochores at meiosis, is consistent with a recent study reporting its comparatively weak kinetochore association in activated gametocytes (Hentzschel et al., 2025), and idea that AKiT7 function may be stage-dependent.

The presence of a minimal machinery that permits transmission efficiency over occasional segregation errors, whilst retaining the capacity for greater fidelity of chromosome segregation when required, aligns with eukaryotic principles observed where the SAC can be modulated or bypassed entirely under certain physiological or developmental constraints. Early *Drosophila* embryos exhibit reduced SAC stringency, facilitating rapid cycling of mitosis without intermittent gap phases (Su et al., 1998; McCleland and O’Farrell, 2008). Similarly, budding yeast tolerates deletion of SAC components *MAD1-3*, increasing chromosome missegregation but without compromising viability under optimal growth conditions (Li and Murray, 1991; Weiss and Winey, 1996).

Permissive, error-prone chromosome segregation in *Plasmodium* may help explain the marked sequence divergence and compositional reduction of its kinetochores compared to other eukaryotes. Weaker error correction would allow centromere-biased mis-attachments (e.g. merotelic or syntelic) to persist, relaxing purifying selection on kinetochore interfaces and accelerating turnover of ‘selfish’ centromere variants. From a therapeutic standpoint, these divergences may represent exploitable vulnerabilities. The stage-specific CPC configurations that safeguard transmission in mosquito stages appear indispensable, yet they are less stringently monitored than canonical SAC pathways. Targeting CPC function genetically or pharmacologically could therefore selectively disrupt parasite proliferation and block transmission, while the essentiality of these processes may limit evolutionary escape routes. Consistent with the *Plasmodium* CPC as an attractive drug target, a recent study has chemically validated the human-infecting parasite *Plasmodium falciparum* ARK1, showing that the human Aurora B inhibitor hesperadin blocks PfARK1 with nanomolar potency across blood, liver, and male gamete stages, providing proof-of-concept that pharmacological targeting of CPC function can deliver multi-stage antimalarial activity (Langeveld et al., 2025).

Taken together, our findings reveal that *Plasmodium* employs a modular CPC to balance propagation and fidelity of chromosome segregation, an evolutionary compromise that exposes vulnerabilities exploitable for transmission-blocking strategies.

### Supporting information

**Table S1. Fidelity of artificial chromosome inheritance across life-cycle stages.** Flow cytometry and microscopy quantification of *Plasmodium* artificial chromosome (PAC) segregation in *P. berghei* male-only lines (fd1-) across 14 intra-erythrocytic cycles and during fertilization. Data include percentages of GFP+BFP+ (centromere-containing PAC), GFP-only (centromere-deficient PACΔcen), total ookinete counts, and inheritance percentages in blood-stage parasites and mature ookinetes.

**Table S2. ARK1 interactome during microgametogenesis.** Mass spectrometry enrichment of proteins co-purifying with ARK1-3xHA in male-only *fd^1-^* gametocytes under native and graded formaldehyde (FA) crosslinking conditions. Relative abundance values are shown as NSAF values alongside AKiT1-3xHA control immunoprecipitates.

**Table S3. Quantification of spindle and MTOC organization during microgametogenesis.** Individual cell measurements for full-cell U-ExM projections of parental and mutant lines, including spindle poles (SP), inner MTOCs (IM), outer MTOCs (OM), over-amplified MTOCs (O), and ratios (e.g. SP/IM, IM/OM).

**Table S4. Mosquito infection outcomes following ARK1 or INCENP1 depletion.** Enumeration of midgut and salivary gland infections per mosquito across biological replicates at 8, 14 and 22 days post-blood-meal.

**Table S5. Meiotic spindle and kinetochore morphology in zygotes/ookinetes.** Individual cell measurements for full-cell U-ExM projections of parental and mutant lines, including nuclear volume, spindle length, ellipticity, DNA content, and normalized spindle parameters.

**Table S6. ARK1 interactome during meiosis and mixed gametocytes.** Mass spectrometry enrichment of proteins co-purifying with ARK1-3xHA in female-only *fd^1-^*gametocytes 8h post-fertilisation (sheet. 1) and mixed *2.34* gametocytes populations 6 minutes post-activation. Relative abundance values are shown as NSAF values alongside AKiT1-3xHA control immunoprecipitates.

**Table S7. Oligonucleotides and genetic constructs used in this study.** Sequences and descriptions of primers for tagging, UTR exchange, and integration, including plasmid backbones and targeted genes.

**Table S8. Antibodies and stains used in this study.** Details of antibodies, fluorescent dyes, and stains including source, dilution, application (IFA, U-ExM, Western blot), and lot numbers.

**Video S1-S7. Representative U-ExM 3D reconstructions.** Full cell projections displaying the methodology used for analysis (S1), parental microgametocyte at 6 minutes post-activation (S2), P_SKA_-ark1 microgametocyte at 6 minutes post-activation (S3), parental ookinete at 8 hours of development (S4), P_CLAG_-ark1 ookinete at 8 hours of development (S5), fully developed parental ookinete at 24 hours (S6), fully developed P_CLAG_-ark1 ookinete at 24 hours of development (S7). Each cell is rotated on its axis and stains shown sequentially for AKiT5-mNG-3xHA (green), DNA (SYTOX), microtubules (α/β-tubulin) and spindle/basal body MTOCs (NHS ester).

## Author Contributions

Conceptualization: L. Brusini. Methodology: L. Brusini, M. Roques, C. Niu. Software: L. Brusini, C. Niu. Formal analysis: L. Brusini, M. Roques, C. Niu. Investigation: L. Brusini, M. Roques, C. Niu. Original draft preparation: L. Brusini. Review and editing: L. Brusini, M. Roques, C. Niu, M. Brochet. Supervision: M. Brochet, V. Heussler. Funding acquisition: L. Brusini, M. Brochet, M. Roques, V. Heussler. All authors contributed to the article and approved the submitted version.

## Acknowledgments

Mass spectrometry and initial quantitation were performed at the Proteomics Core Facility, University of Geneva (unige.ch/medecine/proteomique). We thank in particular the service of Alexandre Hainard and Carla Pasquarello Mosimann in preparation and running of mass spectrometry samples. Microscopy was performed at the Bioimaging core facility, University of Geneva (unige.ch/medecine/bioimaging/en/bioimaging-core-facility), we thank for the technical assistance of Olivier Brun and François Prodon, and the Photonic Bioimaging Center, University of Geneva (unige.ch/bioimaging/photonic/), and we thank for the technical assistance of Christoph Bauer, Jérôme Bosset and Nicolas Liaudet. We thank for the technical assistance of Ruth Rehmann, University of Bern, for mosquito rearing and managing the mouse facility. We thank Oliver Billker, Umeå University, for providing us with *P. berghei* ANKA *fd^1-^* and *md^4-^* lines (Sayers et al., 2024). This work was supported by a Novartis young investigator award #22C170 L. Brusini, University of Geneva maître-assistant salary L. Brusini, Swiss National Science Foundation grant 310030_208151 M. Brochet, Swiss National Science Foundation grant 310030_182465 V. Heussler.

## Data and materials availability

All data needed to evaluate the conclusions in the paper are present in the paper and/or the supplementary materials. Correspondence and requests for materials should be addressed to L. Brusini and M. Brochet.

## Methods

### Ethics statement

All animal experiments were conducted with the authorization number GE283C for University of Geneva and BE118/22 for University of Bern, according to the guidelines and regulations issued by the Swiss Federal Veterinary Office and carried out in strict accordance with the guidelines of the Swiss Tierschutzgesetz (TSchG; Animal Rights Laws).

### Generation of transgenic parasites

The oligonucleotides used to generate transgenic parasite lines are in Table. S7.

For C-terminal tagging of *P. berghei* proteins, constructs were derived from pCPY-0 that insert the coding sequence for mBFP2-2xTy at the C terminus of the target ORF—pCPY-0-mBFP2-2xTy. Briefly, sequences comprising ∼500 bp from the C-terminus of the coding sequence and ∼500 bp from the immediate 3’ UTR for genes ARK1, INCENP1, INCENP2 and AKiT7 were cloned into KpnI and XhoI sites upstream to mBFP2-2xTy coding sequence, along with a NotI linearisation site between the targeting sequences.

For placing *P. berghei* genes under blood-stage expression, constructs were derived from pNP-0 (Brusini et al., 2022), that insert the 5’UTR sequence for the cytoadherence-linked asexual protein, P_CLAG_ (PBANKA_1400600) or spindle and kinetochore associated complex protein SKA2, P_SKA_ (PBANKA_0405800), upstream to a 2xTy coding sequence at the N terminus of the target ORF—pNP-CLAG-2xTy-0 and pNP-SKA-2xTy-0, respectively. Similarly, sequences comprising ∼500 bp from the C-terminus of the coding sequence and ∼500 bp from the immediate 5’ UTR for ARK1, INCENP1 and INCENP2 were cloned into ApaI and EcoRI sites downstream to the 2xTy coding sequence, along with a NotI linearisation site between the targeting sequences.

For generation of *P. berghei* artificial chromosomes (PACs), constructs were newly derived from pBAT1 (Kooij et al., 2012) and PCEN (Iwanaga et al., 2010). Briefly, a 478bp region including the telomeres TelA and TelB was excised from PCEN and inserted into a HindIII site in pBAT1_PB401 to make pBAT1-mCherry_HSP70-GFP-telo. The *P. berghei* centromere 5 region flanked by EcoRV and SacI sites was amplified with MB1700 and MB1701 and inserted downstream to DHFR-FPGS 3’UTR to generate pPAC-mCherry_HSP70-GFP. The gene encoding hDHFR was excised by XhoI and SacI and replaced with one encoding hDHFR and mBFP2 separated by a T2A skip peptide and driven by eEF1α 5’UTR, to generate pPAC-mCherry_HSP70-GFP_eEF1α- hDHFR-T2A-mBFP2. For expression in male gametocytes, the region between BstBI and PvuII sites containing HSP70-GFP was replaced with ORFs for either ARK1 or ark1^K61R^ fused to 2xMyc2xSTREP at the N terminus and flanked by 5’ and 3’ UTRs from PBANKA_1431400, whose transcripts are restricted to male gametocytes (malariacellatlas.org/atlas/pb/), generating pPAC- mCherry_male-ARK1_eEF1α-hDHFR-T2A-mBFP2. For monitoring female gametocytes and ookinetes, mScarlet-I ORF driven by P28 5’UTR replaced mCherry between SalI and SphI sites upstream to PPPK 3’UTR, to generate pPAC-mScI-PPPK_male-ARK1_eEF1α-hDHFR-T2A- mBFP2.

### Plasmodium berghei maintenance and transfection

*P. berghei* ANKA strain-derived clone 2.34 (Billker et al., 2004) together with derived transgenic lines were grown and maintained in CD1 outbred and BALB/c mice. 6-to-8 week-old mice were obtained from Charles River Laboratories and females were used for all experiments. Mice were specific pathogen free (including *Mycoplasma pulmonis*) and subjected to regular pathogen monitoring by sentinel screening. Mice were housed in individually ventilated cages furnished with a cardboard mouse house and Nestlet, maintained at 21 ± 2°C under a 12 h light/dark cycle and given commercially prepared autoclaved dry rodent diet and water ad libitum. The parasitaemia of infected animals was determined by microscopy of methanol-fixed and Giemsa-stained thin blood smears. For gametocyte production, parasites were grown in mice that had been phenyl hydrazine treated three days before infection. Exflagellation was induced in exflagellation medium (RPMI 1640 containing 25 mM HEPES, 4 mM sodium bicarbonate, 5% fetal calf serum (FCS), 100 M xanthurenic acid, pH 7.8). For gametocyte purification, parasites were harvested in suspended animation medium (SA; RPMI 1640 containing 25 mM HEPES, 5% FCS, 4 mM sodium bicarbonate, pH 7.20) and separated from uninfected erythrocytes on a Histodenz/Nycodenz cushion made from 48% of a Histodenz/Nycodenz stock (27.6% [w/v] Histodenz/Nycodenz [Sigma/ Alere Technologies] in 5.0 mM TrisHCl, 3.0 mM KCl, 0.3 mM EDTA, pH 7.20) and 52% SA, final pH 7.2. Gametocytes were harvested from the interface.

Schizonts for transfection were purified from overnight *in vitro* culture on a Histodenz cushion made from 55% of the Histodenz/Nycodenz stock and 45% PBS. Parasites were harvested from the interface and collected by centrifugation at 500 g for 3 min, resuspended in 25 μl Amaxa Basic Parasite Nucleofector solution (Lonza) and added to 10 µg DNA dissolved in 10 µl H_2_O. Cells were electroporated using the FI-115 program of the Amaxa Nucleofector 4D. Transfected parasites were resuspended in 200 μl fresh RBCs and injected intraperitoneally into mice. Parasite selection with 0.07 mg/mL pyrimethamine (Sigma) in the drinking water (pH ∼4.5) was initiated one day after infection.

For mosquito feeding, mice were injected with *P. berghei*-infected erythrocytes or phenylhydrazine hydrochloride (phz, 6 mg/mL in PBS; Sigma Aldrich 114715) by intraperitoneal injection. When parasitemia reached 2-5%, mice were euthanized by CO2 and parasitized blood was isolated by heart puncture and intravenously passaged into pre-treated phz mice. After day 3 post-injection, parasitemia was superior to 10% (gametocytemia > 1%) and mice were ready for mosquito feeding. Microgametocyte activation (exflagellation) was recorded by adding 10-20 µL of a mouse drop blood into 100 µL of ookinete medium (RPMI 1640 containing 25 mM HEPES, 4 mM sodium bicarbonate, 5% fetal calf serum (FCS), 100 µM xanthurenic acid, pH 7.6). The number of exflagellation centers (per field of view: 10-15) was counted 12-15 min post-activation in ookinete medium at room temperature with a phase contrast Nikon microscope using a 60x objective. This mix was then incubated overnight at 20°C and ookinete conversion was recorded the next day on a Leica DM5500 widefield fluorescent microscope using a 100x objective. Mice were anaesthetized with ketamine:xylazine and when mice no longer responded to touch stimulus, were placed on cages with approximately 100-150 mosquitoes. At days 8 and 14 post-mosquito feed, midguts were homogenized to release sporozoites from the oocysts. Ten microliters of oocyst sporozoites suspension was added in a Neubauer counting chamber and all 4 fields were counted to determine the number of midgut sporozoites. At day 22 post-feed, between 7 to 40 mosquito salivary glands were dissected and homogenized to release the sporozoites. 10 µL of sporozoite suspension was added to a Neubauer counting chamber and counted and the average number of salivary gland sporozoites per mosquito calculated.

### Mosquito breeding

Anopheles stephensi mosquitoes were bred at 28 °C, 80% humidity and a 12 h light/dark cycle. Mosquitoes were fed with cotton pads containing 8% fructose and 4-aminobenzoic acid (pABA) (0.2 g/L, Sigma-Aldrich A9878) solution. Female mosquitoes were fed with human blood once a week for egg production. Eggs were kept in water baths supplemented with Liquifry fish food and hatched larvae were grown for 7-12 days until pupae were formed. Larvae were fed with fish food. Pupae were collected in water bowls and placed in mosquito cages for hatching. After feeds on parasite-infected blood, mosquitoes were kept at 20 °C.

### Immunoblotting

The antibodies and dilutions used for immunoblotting are listed in Table. S8.

Immunoblotting was used to confirm presence of tagged proteins. Actively dividing cells were washed in PBS and resuspended at between 1-5 × 10^5^ cells μl^-1^. Lysis was in Laemmli buffer (2% w/v SDS, 0.4 M 2-mercaptoethanol, 10% glycerol, 50 mM Tris-HCl pH 7.2) at 95°C for 5 min. Lysates containing around 5 × 10^6^ cells were separated on 4-20% polyacrylamide gels (Invitrogen) in running buffer (25 mM Tris, 250 mM glycine and 0.1% w/v SDS). Proteins were electrophoretically transferred to 0.45 μm pore-size nitrocellulose membrane at 1.6V cm^-1^ for 14 h in transfer buffer (25 mM Tris, 192 mM glycine, 0.02% w/v SDS and 10-20% methanol). Membranes were blocked in 5% w/v milk powder in TBS-T (20 mM Tris-HCl pH 7.5, 150 mM NaCl, 0.05% Tween-20) for 1 h. Membranes were incubated in primary antibody (Table. S8) in 1% w/v milk in TBS-T for 1 h and washed in TBS-T. Detection was by secondary peroxidase-conjugated antibody (Table. S8), in 1% w/v milk in TBS-T for 1 h. Membranes were washed and detected by chemiluminescence with Western Lightning ECL (PerkinElmer) and exposure to photographic film.

### Protein localisation

The antibodies and stains used for protein localisation are listed in Table. S8.

For localisation of tagged proteins *in vitro* by native fluorescence, cells were mounted in medium (RPMI 1640 containing 25 mM HEPES, 4 mM sodium bicarbonate, 5% fetal calf serum (FCS), 100 µM xanthurenic acid, pH 7.8), and imaged using an inverted Zeiss Axio Observer Z1 microscope fitted with an Axiocam 506 mono 14-bit camera and Plan Apochromat 63x / 1.4 Oil DIC III objective.

For mosquito stages, mosquitoes were aspirated with a small vacuum device and anaesthetized with chloroform. Sleeping mosquitoes were kept on petri dishes on ice. Infected midguts were extracted with forceps and placed on glass slides with 1 µg/mL Hoechst 33342 (Invitrogen, H1399). To record oocyst numbers, images of 20-25 mosquito midguts were acquired with a Leica DM5500 widefield fluorescent microscope using a 10x objective. Live images of oocysts were acquired with a Leica TCS SP8 laser-scanning confocal microscope with a 100x objective. Images were analyzed in Fiji: the function “Find maxima” was used to measure the number of oocysts per midguts, numbers were rectified manually if needed. From the acquired images, oocyst size was then measured for the control and mutant parasite lines. To measure oocyst size, the ellipse icon and “Measure” function was used to measure the area of oocysts. Per experiment, up to 30 oocyst areas were measured for the parent line where possible.

Ultrastructure Expansion Microscopy (U-ExM) followed published protocols (Gambarotto et al., 2019; Bertiaux et al., 2021) with minor adaptations. Briefly, cells were sedimented on poly-D-lysine (Gibco, A38904-01) coverslips (40 μL/coverslip) during 10 minutes at room temperature (RT) with 2% (v/v) methanol-free formaldehyde (Thermo Fisher Scientific, 28906) in PBS for 6 min, followed by 1 minute in methanol at –20 °C and three PBS washes with 100 mM glycine. Coverslips were incubated overnight at 37 °C in a solution of 1.4% (v/v) formaldehyde (Sigma- Aldrich F8775) and 2% (v/v) acrylamide (Sigma-Aldrich, A4058) in PBS prior to gelation in 10% (w/v) APS (Thermo Scientific, 17874) /10% (v/v) TEMED (Thermo Scientific, 17919) /Monomer solution (23% (w/v) sodium acrylate (AK Scientific, R624-100g), 10% (v/v) acrylamide (Sigma-Aldrich, A4058), 0.1% (w/v) N,N′-methylenbisacrylamide (Sigma-Aldrich, M1533), PBS 10X) during 1 hour at 37°C. Gels were denatured at 95°C for 1.5 hours in denaturation buffer (200 mM SDS (PanReac AppliChem, A3942), 200 mM NaCl, 50 mM Tris, pH 9). Gels were expanded in ddH2O overnight in three consecutive water incubations. The following day, gels were cryopreserved unstained after the first expansion round by washing 3× in 50% (v/v) glycerol (Sigma-Aldrich, 49767) in Milli-Q water and stored at –20 °C. On staining day, gels were washed in PBS and blocked in 3% BSA (PanReac AppliChem, A1391) in PBS for 30 min at 37 °C. Primary antibodies were diluted in 2% BSA/PBS and incubated for 3 h at 37 °C. Gels were washed in PBS-Tween 0.1% prior incubation with secondary antibodies (Table. S8) for 2.5 hours at 37°C. After washing in PBS-Tween 0.1%, gels were incubated for 1 h at room temperature with NHS ester and SYTOX both diluted in PBS. Gels were then washed in PBS-T and re-expanded in ddH2O overnight in three consecutive water incubations. The expansion factor (between 4.2 and 4.6) was determined by measuring the post-expansion gel diameter and comparing it to the initial 12 mm coverslip size. Confocal imaging was performed using either a Leica TCS SP8 with HC PL APO CS2 100×/1.40 oil objective and LAS X 3.5.7.23225 or a Leica Stellaris 8 FALCON with HC PL APO CS2 63×/1.20 water objective and LAS X 4.8.1.29271. Deconvolution was performed using Huygens Essential 24.04 (Scientific Volume Imaging B.V.), with identical parameters applied across samples. For each condition, 17–30 cells were imaged (z-stacks) under consistent acquisition settings. All images of fluorescent proteins were captured at room temperature. Images were viewed with Fiji (Schindelin et al., 2012) and volume and distance data were obtained using Imaris 9.5 (Bitplane). NHS ester 405-stained basal bodies (MTOCs) were detected with Spot objects. HA-tagged AKiT1, stained with Alexa Fluor 488, were detected with Surface objects. Kinetochore volumes were obtained from the Surface objects. Axonemes and spindle microtubules were labeled with α+β tubulin antibodies and detected with goat anti-guinea pig Alexa Fluor 568. For the 6 and 12 minutes conditions, male gametocytes were identified by their axonemes during acquisition. Spindle poles were manually categorized as segregated or non-segregated. Defective spindles were manually counted. In the 8h zygotes, the spindle length was measured as the distance between two MTOCs. In the 24 h ookinetes, DNA stained with SYTOX Deep Red was additionally used for quantification of nuclear volumes with Surface detection. Statistics were computed using R (http://r-project.org), applying the measured expansion factor to infer non-expanded dimensions.

### Flow cytometry

To measure the fluorescence ratio between *P. berghei* populations from the infected mice, a few drops of blood from the tail vein of infected mice were added to 500 μL of 1 x PBS and measured by flow cytometry using the MoFlo ASTRIOS EQ, as described (Dekel et al., 2017). mCherry (excitation 569–593), GFP (excitation 488–507) and BFP (excitation 383–448) lasers were used to select mScarlet-I-, GFP- and BFP-expressing transgenic parasites, respectively. The data were analyzed using the FCS Express Flow Cytometry Analysis Software (version 7.22.0006).

### Chromosome loss assay

To assess the fidelity of chromosome segregation across parasite life-cycle stages, we developed a *Plasmodium artificial chromosome* assay (Fig. S1B). PACs encoding a centromeric region from *P. berghei* chromosome 5 flanked by telomeric repeats, together with two independent fluorescent markers: (i) mBFP2 expressed constitutively from the eEF1α promoter to report inheritance of the PAC in asexual blood stages, and (ii) mScarlet-I expressed from the female/ookinete-specific P28 promoter to report inheritance following fertilisation and ookinete development. A control construct lacking the centromeric region (PACΔcen) was used to distinguish centromere-dependent from random segregation.

For asexual stage assays, *fd1-::PAC* parasites were serially passaged through 14 rounds of intra- erythrocytic division in mice. Blood was collected at defined time points, and PAC inheritance was quantified by flow cytometry as the proportion of GFP+ BFP+ cells among the total infected population, compared to background levels in PACΔcen parasites. Fidelity per schizogeny cycle was calculated assuming cumulative segregation errors accumulate multiplicatively across cycles, using the formula:

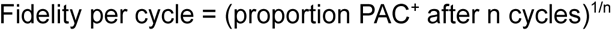

For sexual stage assays, male-only *fd^1-^::PAC* parasites were crossed with untransformed female- only *md^4-^* parasites, and zygotes were cultured to ookinetes *in vitro*. PAC inheritance was scored by fluorescence microscopy as the proportion of ookinetes expressing mScI relative to the total number of ookinetes imaged at 48 hours post-fertilisation. In all cases, values are reported as mean percentages from at least three independent experiments, with thresholds for background determined from controls.

### Immunopurification

For the purification of aurora kinase complexes, gametocytes and zygotes were purified from 5- 10 ml *P. berghei* infected mouse blood. Cells were washed twice in HKMEG (150 mM KCl, 150 mM glucose, 25 mM HEPES, 4 mM MgCl2, 1 mM EGTA, pH7.8) containing 100 x protease inhibitor cocktail and 20 μM MG132. Cells were treated with 0% (“no”), 0.1% (“low”), or 1% (“high”) formaldehyde in HKMEG for 10 min, quenched with 10 ml of 1 M glycine, and lysed in HKMEG containing 1% (vol/vol) Nonidet P-40, 0.5% sodium deoxycholate, 0.1% SDS, 1mM dithiothreitol (DTT), 100 x protease inhibitor cocktail and 20 μM MG132. Lysate was sonicated in an ultrasonic water bath for 20 min applied for 50% of the cycle and cleared by centrifugation at >20,000 g for 30 min. The soluble fraction was then allowed to bind to affinity-purified rat anti-HA antibody (Sigma) attached to paramagnetic beads (Dynabeads protein G; Invitrogen) for 4 hours. Beads were washed in HKMEG containing 0.1% (vol/vol) Nonidet P-40, 0.5mM dithiothreitol. Beads were re-suspended in 100 µl of 6 M urea in 50 mM ammonium bicarbonate (AB). 2 µl of 50 mM dithioerythritol (DTE) were added and the reduction was carried out at 37°C for 1 hr. Alkylation was performed by adding 2 µl of 400 mM iodoacetamide for 1 hr at room temperature in the dark. Urea was reduced to 1 M by addition of 500 ml AB and overnight digestion was performed at 37°C with 5 ml of freshly prepared 0.2 mg/ml trypsin (Promega) in AB. Supernatants were collected and completely dried under speed-vacuum. Samples were then desalted with a C18 microspin column (Harvard Apparatus) according to manufacturer’s instructions, completely dried under speed-vacuum and stored at -20°C.

### Mass spectrometry

For protein identification, samples were diluted in 20 µl loading buffer (5% acetonitrile, 0.1% formic acid [FA]) and 2 µl were injected onto the column. LC-ESI-MS/MS was performed either on a Q-Exactive Plus Hybrid Quadrupole-Orbitrap Mass Spectrometer (Thermo Fisher Scientific) equipped with an Easy nLC 1000 liquid chromatography system (Thermo Fisher Scientific) or an Orbitrap Fusion Lumos Tribrid mass Spectrometer (Thermo Fisher Scientific) equipped with an Easy nLC 1200 liquid chromatography system (Thermo Fisher Scientific). Peptides were trapped on an Acclaim PepMap 100, 3 µm C18, 75 µm x 20 mm nano trap-column (Thermo Fisher Scientific) and separated on a 75 µm x 250 mm (Q-Exactive) or 500 mm (Orbitrap Fusion Lumos), 2 µm C18, 100 Å Easy-Spray column (Thermo Fisher Scientific). The analytical separation used a gradient of H_2_O/0.1% FA (solvent A) and CH_3_CN/0.1% FA (solvent B). The gradient was run as follows: 0 to 5 min 95% A and 5% B, then to 65% A and 35% B for 60 min, then to 10% A and 90% B for 10 min and finally for 15 min at 10% A and 90% B. Flow rate was 250 nL/min for a total run time of 90 min. Data-dependent analysis (DDA) was performed on the Q-Exactive Plus with MS1 full scan at a resolution of 70,000 Full width at half maximum (FWHM) followed by MS2 scans on up to 15 selected precursors. MS1 was performed with an AGC target of 3 x 10^6^, a maximum injection time of 100 ms and a scan range from 400 to 2000 m/z. MS2 was performed at a resolution of 17,500 FWHM with an automatic gain control (AGC) target at 1 x 10^5^ and a maximum injection time of 50 ms. Isolation window was set at 1.6 m/z and 27% normalised collision energy was used for higher-energy collisional dissociation (HCD). DDA was performed on the Orbitrap Fusion Lumos with MS1 full scan at a resolution of 120,000 FWHM followed by as many subsequent MS2 scans on selected precursors as possible within a 3 s maximum cycle time. MS1 was performed in the Orbitrap with an AGC target of 4 x 10^5^, a maximum injection time of 50 ms and a scan range from 400 to 2000 m/z. MS2 was performed in the Ion Trap with a rapid scan rate, an AGC target of 1 x 10^4^ and a maximum injection time of 35 ms. Isolation window was set at 1.2 m/z and 30% normalised collision energy was used for HCD.

### Database search

Raw data were processed using Proteome Discoverer 2.3 software (Thermo Fisher Scientific). Briefly, spectra were extracted and searched against the *P. berghei* ANKA database (PlasmoDB.org, release 46) combined with an in-house database of common contaminants using Mascot (Matrix Science, London, UK; version 2.5.1). Trypsin was selected as the enzyme, with one potential missed cleavage. Precursor ion tolerance was set to 10 ppm and fragment ion tolerance to 0.02 Da. Carbamidomethyl of cysteine (+57.021) and on peptide N-termini were specified as fixed modifications. Oxidation of methionine (+15.995) as well as phosphorylated serine, threonine and tyrosine were set as variable modifications. The search results were validated with a Target Decoy PSM validator. PSM and peptides were filtered with a FDR of 1%, and then grouped to proteins, again with a FDR of 1% and using only peptides with high confidence level. Both unique and razor peptides were used for quantitation and protein and peptide abundances were based on signal-to-noise (S/N) values of reporter ions. Abundances were normalised on ‘Total Peptide Amount’ and then scaled with ‘On Controls Average’ (i.e. using the reference sample channel). All the protein ratios were calculated from the medians of the summed abundances of replicate groups and associated p-values were calculated with an ANOVA test based on individual protein or peptides. GO term enrichment was performed using PlasmoDB.

Enrichment analyses were performed in the statistical programming package ‘R’ (r-project.org). Quantitative values were analyzed as NSAF value versus control immuno-purification.

---

**Figure S1.**
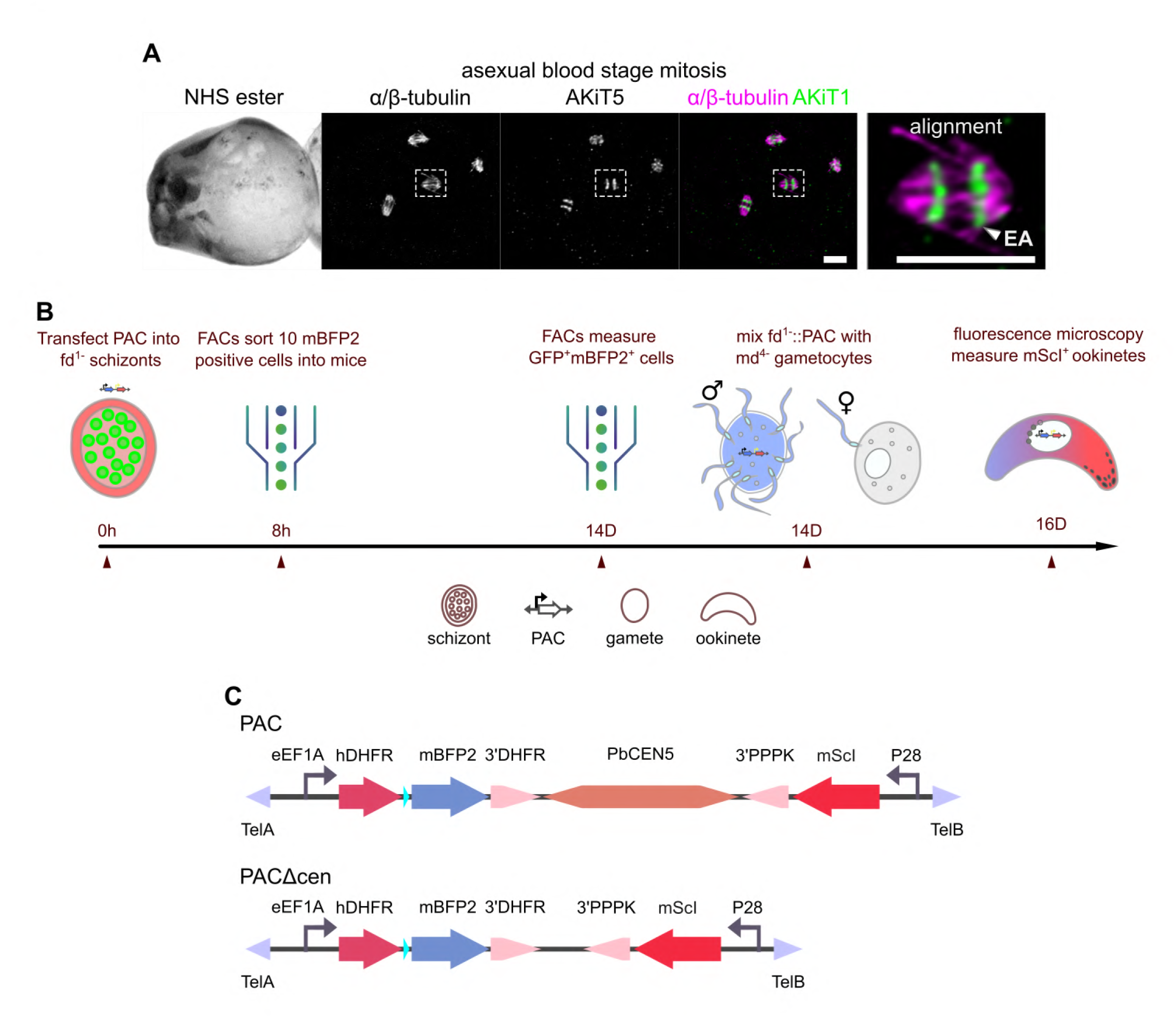
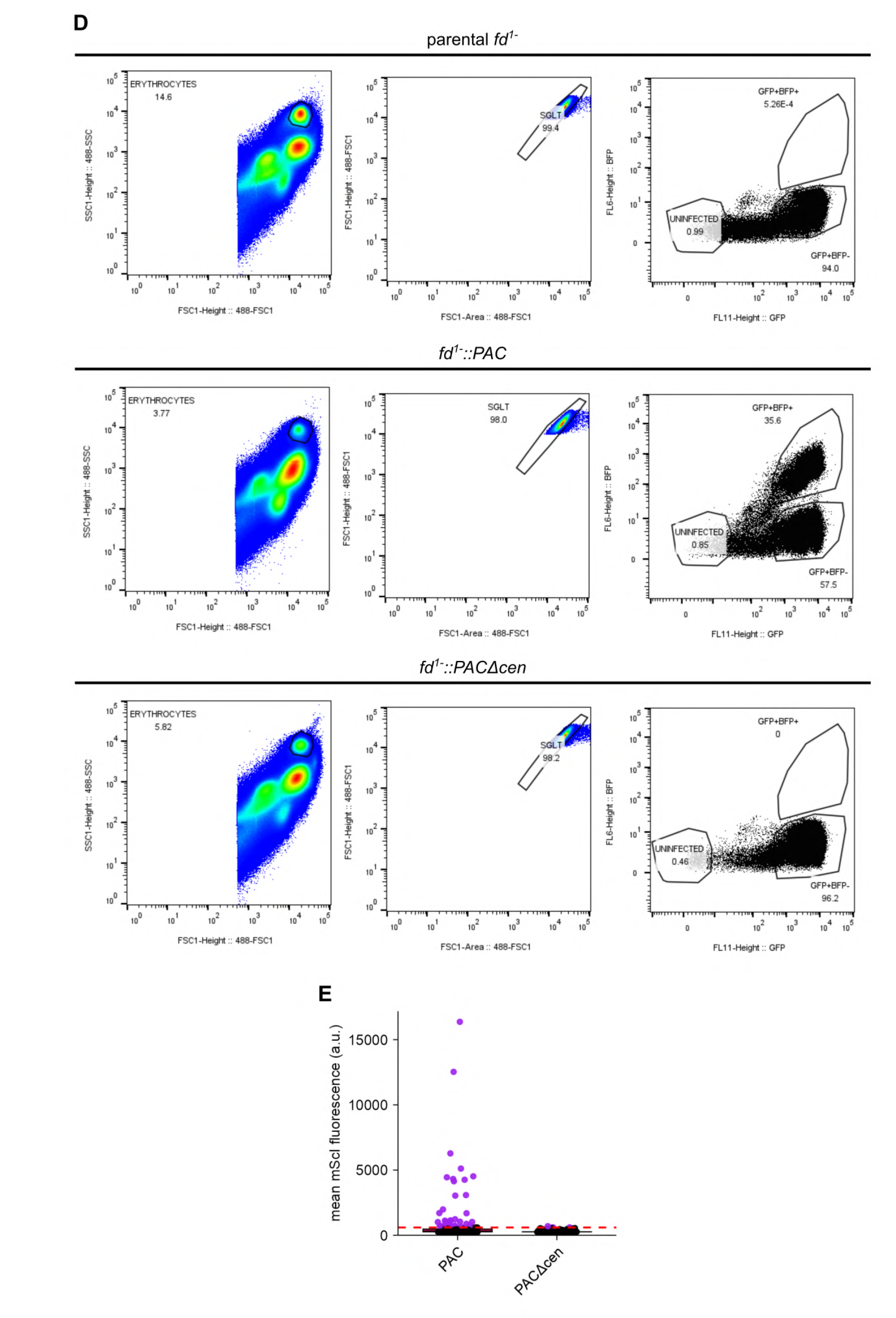
Chromosome segregation analysis between asexual and sexual stage *P. berghei*. **(A)** U-ExM micrographs of intra-erythrocytic asexual *Plasmodium berghei* expressing the tagged kinetochore component AKiT5-mNG-3xHA (green). Counter-staining of the spindle (α/β-tubulin) and spindle/basal body MTOCs (NHS ester) also shown. Representative panels displaying alignment of kinetochores at the nuclear equator and end-on kinetochore attachment (EA). Scale bar, 1 μm (non-expanded). (B) Workflow used to measure the transmission of *Plasmodium* artificial chromosomes (PAC) between asexual and sexual stage cells. (C) Schematic representation of engineered PACs either with or without the centromeric region from chromosome 5, in addition to gene cassettes for the expression of mBFP2 and mScarlet-I and flanked by telomeric DNA. (D) Flow cytometric analysis of PAC positive (GFP^+^BFP^+^) cells following 14 serial passages in mice, compared to parental and PAC lacking the centromere (PACΔcen). (E) Fluorescence microscopy analysis of PAC positive (mScI^+^) ookinetes 48 hours post-fertilisation of female-only *md^4-^* gametocytes by *fd^1-^::PAC* male-only gametocytes. Red dotted line represents 99% quantile used to determine mScI^+^ background fluorescence.

**Figure S2.**
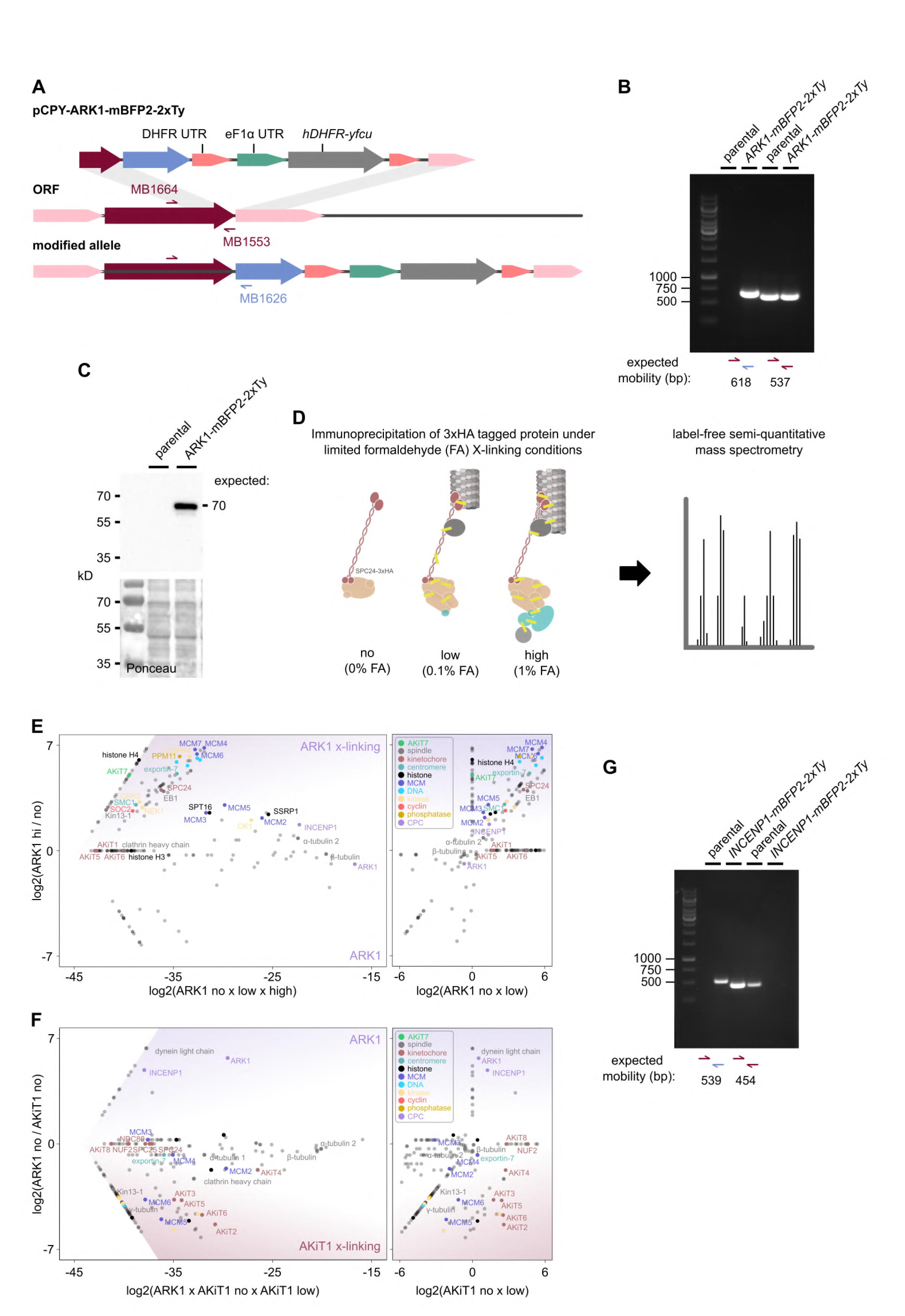
Generation of *ARK1-mBFP2-2xTy P. berghei* line and immunoprecipitation of ARK1 and kinetochore components. (A&B) Genomic DNA integration PCR from *P. berghei* tagged with mBFP2-2xTy at the C-terminus of endogenous *ARK1* and alongside parental controls. (C) Immunoblots of ARK1-mBFP2-2xTy expressing parasites compared to parental controls, probed with monoclonal α-HA antibodies. Protein loading is shown by Ponceau S stain. (D) General workflow for immunoprecipitation under limited cross-linking and mass spectrometry. (E) Relative enrichment of proteins identified by mass spectrometry following immunoprecipitation of ARK1-3xHA (magenta) under native and cross-linking conditions, and AKiT1-3xHA (burgundy). ARK1 and AKiT1-interacting proteins highlighted. Intensities of proteins not detected for either immunoprecipitation set to arbitrary minimum value.

**Figure S3.**
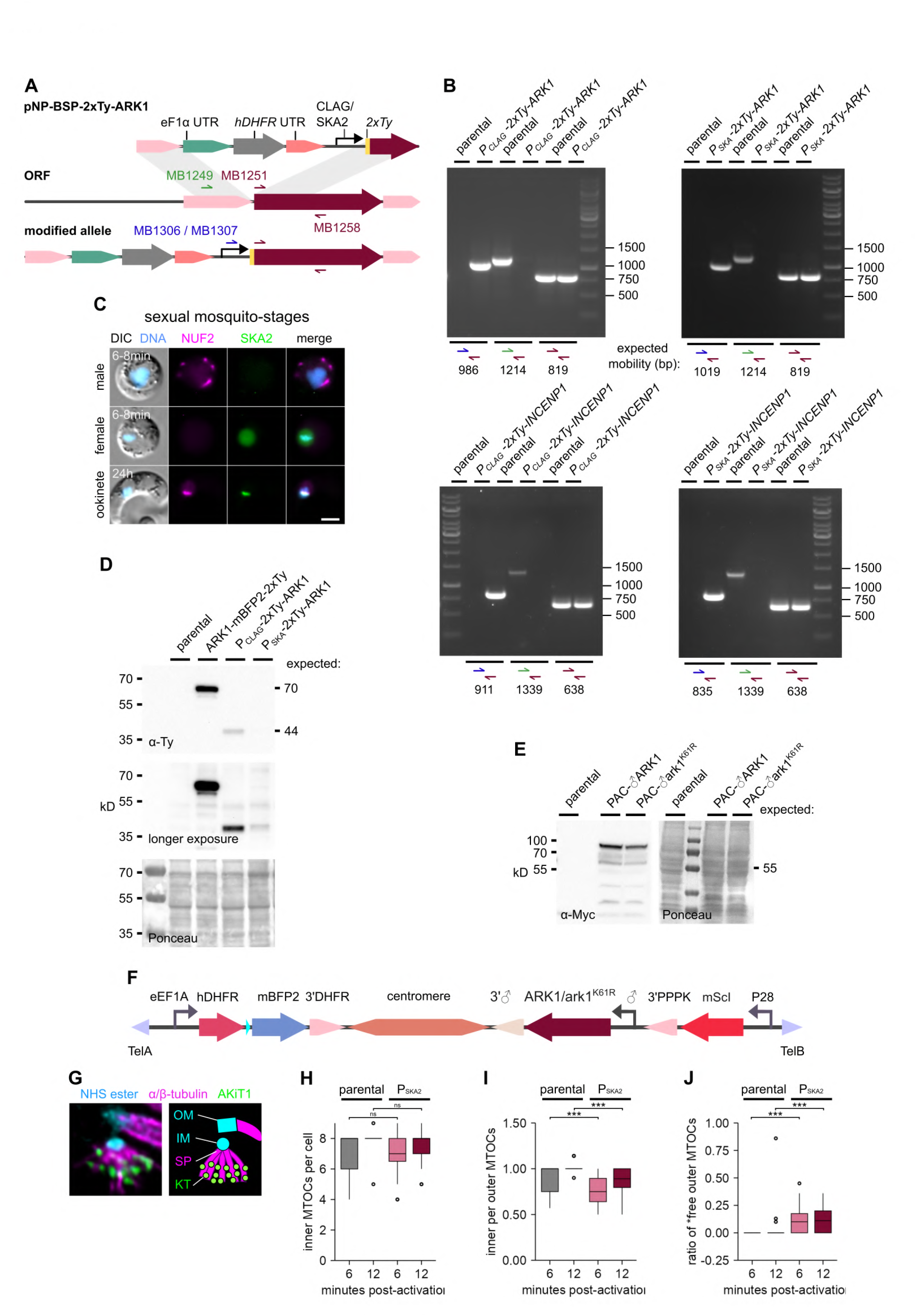
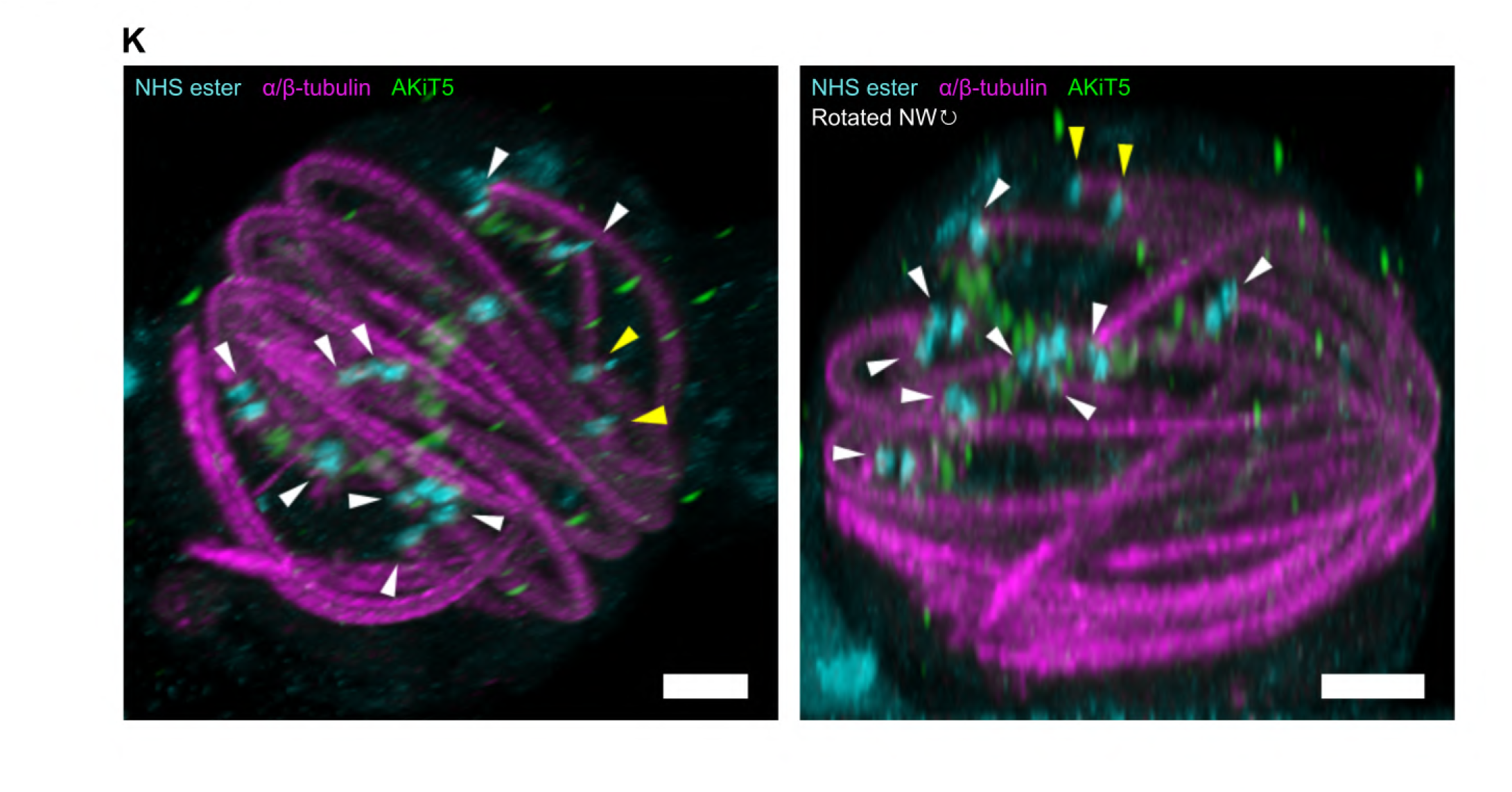
Depletion of ARK1 and INCENP1 impairs proper spindle and kinetochore dynamics during microgametogenesis. (A&B) PCR on genomic DNA of *P. berghei* with integrated CLAG or SKA2 promoters and tagged with 2xTy at the N-terminus of endogenous *ARK1* and *INCENP1*, and alongside parental controls. (C) Image taken from Brusini et al., 2022; Fig. 1B. Micrographs of live native fluorescence in *P. berghei* mosquito stages expressing tagged kinetochore components NUF2-mScarlet-I (magenta) and SKA2-mNG-3xHA (green). Counter- staining of DNA with Hoechst 33342 (cyan) and differential interference contrast images are also shown. Scale bar, 2 μm. (D) Immunoblots of proteins from *P. berghei* gametocytes expressing *2xTy-ARK1* under the regulation of P_CLAG_ or P_SKA_ promoters reveal a marked decrease in protein levels compared to endogenous ARK1-mBFP2-2xTy, and alongside parental controls. Probed with mouse hybridoma α-Ty antibodies, protein loading is shown by Ponceau S stain. (E) Immunoblots from *P. berghei* gametocytes expressing ectopic *2xMyc2xSTREP-ARK1* under the regulation of the male (♂) promoter from PBANKA_1431400 (Malaria Cell Atlas; malariacellatlas.org/) and alongside parental controls. Probed with mouse hybridoma α-Myc antibodies, protein loading is shown by Ponceau S stain. (F) Schematic representation of engineered PACs encoding the centromeric region from chromosome 5, and cassettes for the expression of *ARK1* or *ark1^K61R^*under the regulation of the male (♂) promoter from PBANKA_1431400, in addition to the expression of mBFP2 and mScI, and flanked by telomeric DNA. (G) Representative micrograph and annotated schematic displaying organelles analyzed by U-ExM, including the inner and outer MTOCs (IM/OM; NHS ester, cyan), spindle pole (SP; α/β- tubulin, magenta) and kinetochores (KT; AKiT5-mNG-3xHA, green). U-ExM of microgametocytes revealed no difference in the number of inner MTOCs (H) in ARK1-depleted cells, however an over-amplification of outer MTOCs (I) and number of “free” outer MTOCs not connected to a spindle apparatus (J). Boxplots show the median (line), interquartile range (box), whiskers (1.5 × IQR), and outliers (points), Wilcoxon tests; ns ≥ 0.05; * < 0.05; ** < 0.01; *** < 0.001. (K) Representative U-ExM micrograph of ARK1-depleted microgametocyte displaying >8 outer MTOCs (arrows) and not attached to inner MTOCs (yellow arrows). Right panel represents north- western rotation of panel on the left. MTOCs (NHS ester), mitotic spindles (α/β-tubulin), DNA (SYTOX) and kinetochores (AKiT5-mNG-3xHA) are shown. Scale Bar: 1 μm (non-expanded).

**Figure S4.**
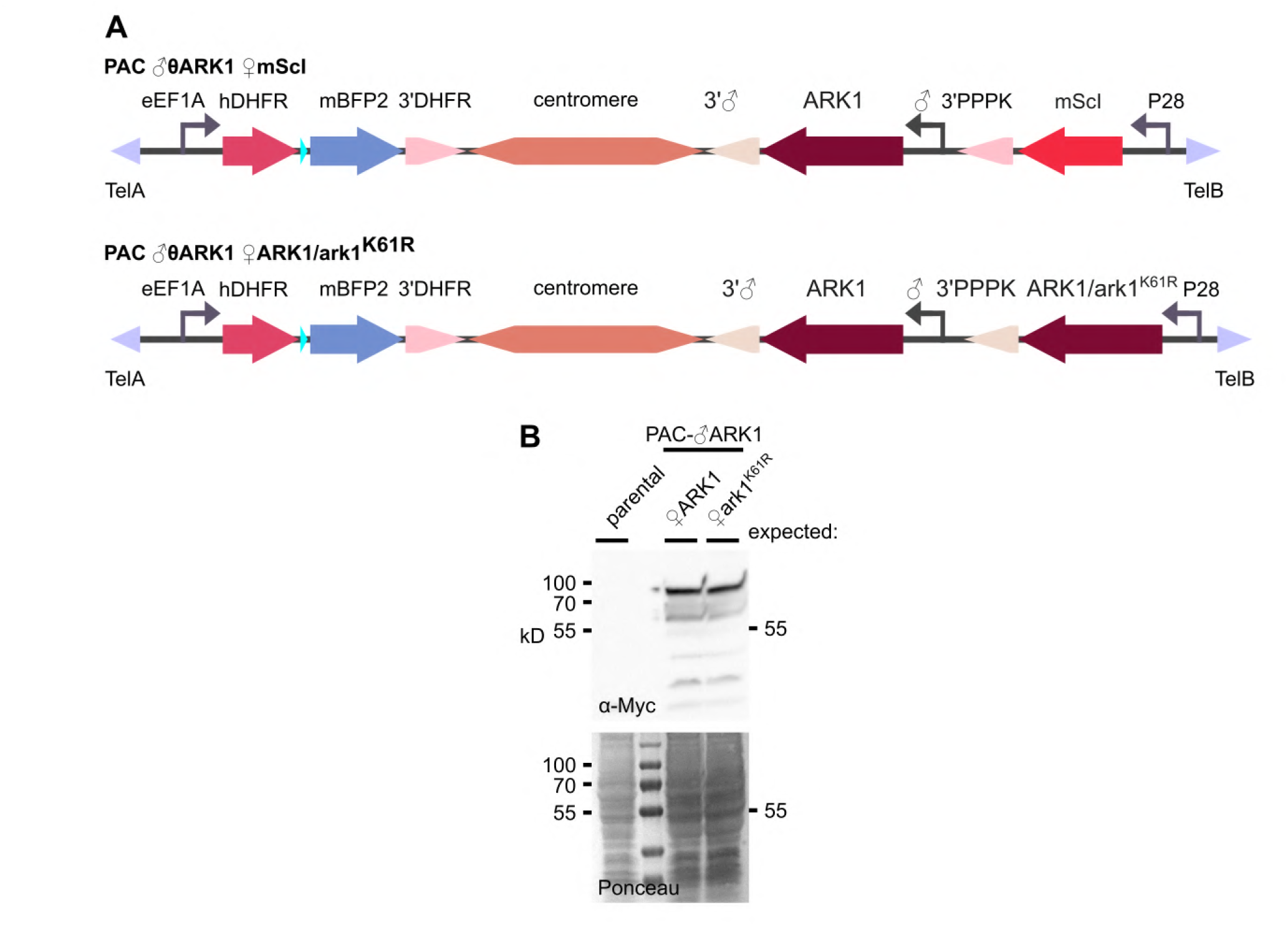
Ectopic PAC expression of ARK1 and ark1^K61R^ in gametocytes and ookinetes. (A) Schematic representation of engineered PACs encoding the centromeric region from chromosome 5, and cassettes for the expression of *ARK1* under the regulation of the male (♂) promoter from PBANKA_1431400, the expression of either mScI, *ARK1* or *ark1^K61R^* under the regulation of the P28 (♀) promoter, in addition to mBFP2, and flanked by telomeric DNA. (B) Immunoblots from *P. berghei* ookinetes expressing ectopic *2xMyc2xSTREP-ARK1* under the regulation of the male (♂) promoter from PBANKA_1431400 and 2*xMyc2xSTREP-ARK1 / ark1^K61R^* from the the P28 (♀) promoter and alongside parental controls. Probed with mouse hybridoma α-Myc antibodies, protein loading is shown by Ponceau S stain.

**Figure S5.**
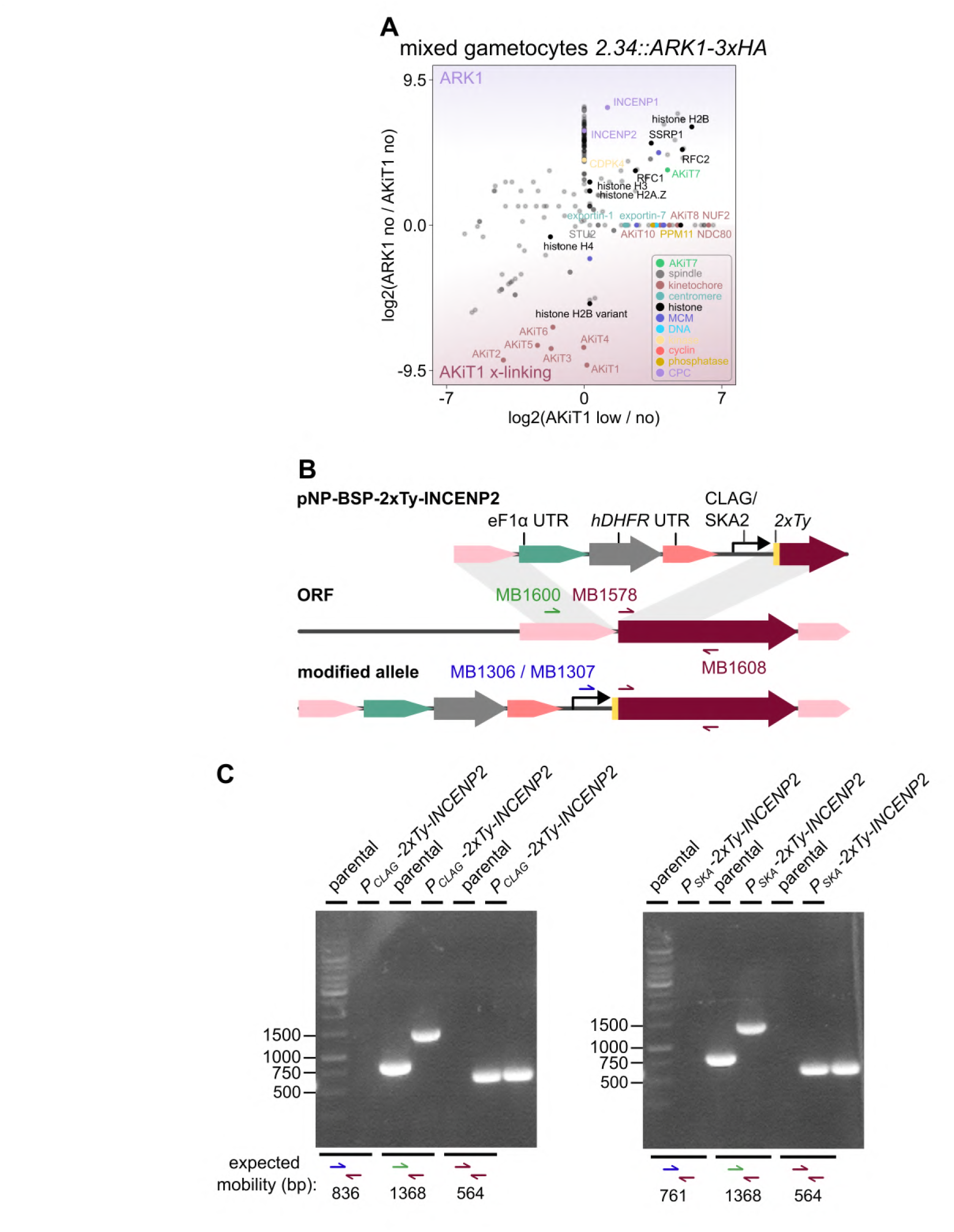
Generation of *P_CLAG_/P_SKA_-2xTy-incenp2 P. berghei*. (A) Relative enrichment of proteins identified by mass spectrometry following immunoprecipitation of ARK1-3xHA (magenta) and AKiT1-3xHA (burgundy) from mixed male and female *2.34 P. berghei* gametocytes 6 minutes post-activation. ARK1 and AKiT1-interacting proteins highlighted. Intensities of proteins not detected for either immunoprecipitation set to arbitrary minimum value. Intensities for all 211 proteins detected are presented in Table. S6. (B&C) Genomic DNA integration PCR from *P. berghei* with integrated CLAG or SKA promoters and tagged with 2xTy at the N-terminus of endogenous *INCENP2* and alongside parental controls.

